# Total evidence tip-dating phylogeny of platyrrhine primates and 27 well-justified fossil calibrations for primate divergences

**DOI:** 10.1101/2021.10.21.465342

**Authors:** Dorien de Vries, Robin M. D. Beck

## Abstract

Phylogenies with estimates of divergence times are essential for investigating many evolutionary questions. In principle, “tip-dating” is arguably the most appropriate approach, with fossil and extant taxa analyzed together in a single analysis, and topology and divergence times estimated simultaneously. However, “node-dating” (as used in many molecular clock analyses), in which fossil evidence is used to calibrate the age of particular nodes a priori, will probably remain the dominant approach, due to various issues with analyzing morphological and molecular data together. Tip-dating may nevertheless play a key role in robustly identifying fossil taxa that can be used to inform node-dating calibrations. Here, we present tip-dating analyses of platyrrhine primates (so-called “New World monkeys”) based on a total evidence dataset of 418 morphological characters and 10.2 kb of DNA sequence data from 17 nuclear genes, combined from previous studies. The resultant analyses support a late Oligocene or early Miocene age for crown Platyrrhini (composite age estimate: 20.7-28.2 Ma). Other key findings include placement of the early Miocene putative cebid *Panamacebus* outside crown Platyrrhini, equivocal support for *Proteropithecia* being a pitheciine, and support for a clade comprising three subfossil platyrrhines from the Caribbean (*Xenothrix*, *Antillothrix* and *Paralouatta*), related to Callicebinae. Based on these results and the available literature, we provide a list of 27 well-justified node calibrations for primate divergences, following best practices: 17 within Haplorhini, five within Strepsirrhini, one for crown Primates, and four for deeper divergences within Euarchontoglires. In each case, we provide a hard minimum bound, and for 23 of these we also provide a soft maximum bound and a suggested prior distribution. For each calibrated node, we provide the age of the oldest fossil of each daughter lineage that descends from it, which allows use of the “CladeAge” method for specifying priors on node ages.

## 1. Introduction

Phylogenies that provide estimates of divergence times between different lineages of organisms (“timetrees”) have the potential to be extremely useful for answering a wide range of questions relating to evolutionary patterns and processes (Blair Hedges and Kumar, 2009; Ho, 2021). There has been a long history of applying such approaches to primates, for example to identify when humans diverged from their closest living relatives (Sarich and Wilson, 1967; Hasegawa et al., 1985; Scally and Durbin, 2012; Schrago and Voloch, 2013; Püschel et al., 2021), to determine when key phenotypic changes (e.g. increases in brain size; Ni et al., 2019; Püschel et al., 2021) occurred, to clarify the timing of significant biogeographical events (e.g. the dispersal of primates to Madagascar and to South America during the Cenozoic; Poux et al., 2005, 2006; Gunnell et al., 2018; Seiffert et al., 2020), and to infer the likely impact of major environmental changes (e.g., shifts in global climate) on primate diversification (Springer et al., 2012; Herrera, 2017; Godfrey et al., 2020).

Calculation of absolute divergence times between lineages usually requires some form of calibration by incorporating known temporal information, and for “deep time” divergences this typically means evidence from the fossil record (Nguyen and Ho, 2020). In a phylogenetic context, arguably the most appropriate approach (at least in principle) is “tip-dating”, in which fossil taxa of known age are incorporated as terminals (“tips”) in the phylogenetic analysis, and this temporal information is used to estimate divergence times at the nodes (Pyron, 2011; Ronquist et al., 2012a; O’Reilly et al., 2015; Zhang et al., 2016; Lee, 2020); this approach, in which fossil and extant taxa can be analyzed together, has the advantage that topology and divergence times are estimated simultaneously, in the context of a single analysis, and so the relationships of the fossil taxa do not need to be assumed a priori.

However, it seems unlikely that large quantities of molecular data will ever become available for fossil taxa older than a few million years old (Allentoft et al., 2012; Millar and Lambert, 2013; Welker, 2018), and so tip-dating analyses that include fossil taxa will typically require morphological data to inform their evolutionary relationships (although note that tip-dating can be used to infer divergence times among fossil taxa based purely on their stratigraphic age if their relationships are constrained a priori; Silvestro et al., 2019b; Button and Zanno, 2020; Lloyd and Slater, 2021). One potential problem with this approach concerns models of morphological character evolution, which are still in their infancy; in particular, the most widely used model for analyzing discrete morphological data (the “Mk” model of Lewis, 2001) makes a number of simplifying assumptions that seem biologically unrealistic (O’Reilly et al., 2015; Wright et al., 2016; Pyron, 2017; Billet and Bardin, 2019; Wright, 2019). However, a simulation study by Klopfstein et al. (2019) suggests that this may not be particularly problematic for the inference of divergence dates, at least when relevant fossil taxa are densely sampled.

More serious is the fact that, even for primates (perhaps the most well-studied of any clade of organisms), morphological datasets are still small in comparison to equivalent molecular datasets, in terms of both taxon and character sampling (Guillerme and Cooper, 2016a). Species-level molecular phylogenies have been available for extant primates for a number of years (e.g. Chatterjee et al., 2009; Perelman et al., 2011; Springer et al., 2012; dos Reis et al., 2018), and population- and subpopulation-level phylogenies will undoubtedly be generated with increasing rapidity. By contrast, even in the most recent morphological datasets for primate phylogenetics (at least, those focused on higher level relationships, e.g., between families), extant taxa are typically scored at the genus level, or only a single species-level representative of each genus is included (e.g. Rasmussen et al., 2019; Gilbert et al., 2020; Seiffert et al., 2020).

Given that scoring of morphological datasets requires anatomical expertise, considerable amounts of time, and access to widely scattered museum collections or high quality (ideally 3D) images of specimens, it is unlikely that future morphological datasets for primates will ever be sampled as densely in terms of taxa as will future molecular datasets. Thus, directly combining a phylogenomic dataset that samples extant primates at the population or subpopulation level with a compatible morphological dataset will result in a total evidence dataset in which most extant terminals will lack morphological data (Guillerme and Cooper, 2016b). Relationships between fossil taxa represented solely by morphological data and extant taxa represented solely by molecular data cannot be resolved without the use of topological constraints, because they do not share any character data in common (Springer et al., 2004; Guillerme and Cooper, 2016b; Hunt and Slater, 2016). In addition, phylogenomic datasets for primates can comprise millions of base pairs of sequence data (e.g. Jameson et al., 2011; dos Reis et al., 2018; Vanderpool et al., 2020), while primate morphological datasets typically comprise a few hundred characters (e.g. Rasmussen et al., 2019; Gilbert et al., 2020; Seiffert et al., 2020), and so combining such datasets will mean that fossil taxa (without molecular data) might end up with >99.99% missing data. The impact of such large amounts of missing data on phylogenetic inference is not fully understood, and may not be as severe as might be expected (Wiens et al., 2005; Guillerme and Cooper, 2016b), but nevertheless can still negatively affect accurate inference of topology, support values, and (crucially for the estimation of divergence times) branch lengths (Lemmon et al., 2009; Simmons, 2012, 2014; Xia, 2014).

For these and other reasons (including, we suspect, the relative unfamiliarity to many bioinformaticians of morphological data and of appropriate methods for analysing it), it seems likely that molecular clock analyses, which use molecular data only, will remain the most common approach for inferring divergence times among lineages. In such analyses, fossil taxa for which molecular data is unavailable cannot be included directly as terminals/tips, but instead can be used to calibrate the age of one or more nodes in the phylogeny (“node-dating”; Nguyen and Ho, 2020). Because fossils are assigned a priori to particular nodes under this approach, it is crucial that each calibrating fossil is assigned appropriately, based on the most accurate and up-to-date information (Parham et al., 2012; Marjanović, 2021).

However, as discussed at length by Marjanović (2021), identifying appropriate fossil calibrations is not trivial, and even comparatively recent lists of calibrations (e.g. Benton et al., 2015) now appear out of date. Such issues affect molecular clock analyses of primates. For example, several studies (Meredith et al., 2011; Perelman et al., 2011; Springer et al., 2012) have used the late Eocene fossil primate *Saharagalago*, which was originally described as a galagid (Seiffert et al., 2003), to place a minimum bound on the age of crown Lorisiformes (= the Galagidae-Lorisidae split). However, a number of subsequent studies - notably, several tip-dating analyses (Herrera and Dávalos, 2016; Gunnell et al., 2018; Seiffert et al., 2020) - have instead placed *Saharagalago* as a stem lorisiform (see summaries in López-Torres and Silcox, 2020; López-Torres et al., 2020), suggesting that it is unsuitable for calibrating the Galagidae-Lorisidae split. Phillips (2016) and Phillips and Fruciano (2018) also found that use of *Saharagalago* for this calibration results in a seeming mismatch in molecular rates, further suggesting this fossil taxon is not a crown lorisiform. Tip-dating may therefore still play a key role in determining divergence times within primates, as part of a two-stage process: tip-dating analyses of smaller (in terms of both taxa and characters) datasets, which ideally use combined morphological and molecular (=total evidence) data and which show good overlap of characters between taxa, can be used to robustly identify fossil taxa suitable for calibrating particular nodes; these calibrations can then be applied to clock analyses of larger, molecular-only datasets.

The way that fossil calibrations are specified in a node-dating analysis is known to have a major impact on the estimated divergence times (Warnock et al., 2012, 2015). Although fossil calibrations have been used to specify normal distributions on the ages of nodes in some studies (e.g. Perelman et al., 2011), it seems more appropriate to view the minimum age of a calibrating fossil as providing a minimum bound on the age of that node (Benton and Donoghue, 2007; Benton et al., 2009; Ho and Phillips, 2009; Parham et al., 2012). Wherever possible, it is appropriate to also specify a maximum bound, otherwise there is no direct constraint on the maximum age of a calibrated node (Phillips, 2016; Marjanović, 2021). If maximum and minimum bounds are specified, then a prior probability distribution between these bounds (and, if these bounds are “soft”, outside them as well) also needs to be specified (Ho and Phillips, 2009). In principle, any distribution could be used, but from a paleontological perspective, perhaps the most defensible are: 1) a uniform distribution, in which there is an equal probability that the divergence occurred at any time between the minimum and maximum bounds, and which appears most appropriate in cases where the fossil record is known or suspected to be very incomplete, such that the oldest calibrating fossil may in fact substantially postdate the age of the divergence being calibrated; 2) an exponential distribution, in which the probability that the divergence is older than the minimum bound decreases exponentially, and which appears most appropriate in cases where the calibrating fossil is suspected to be very close in time to the actual time of divergences (Ho and Phillips, 2009).

Analytical methods for inferring maximum bounds and prior probability distributions on calibrations have been proposed (Marshall, 2008; Wilkinson et al., 2011; Nowak et al., 2013; Matschiner et al., 2017; Matschiner, 2019), but these often require estimates of parameters such as diversification and/or sampling rates that are not always easy to obtain, even for primates (but see Silvestro et al., 2014, 2019a; Herrera, 2017). For this reason, maximum bounds are typically based on somewhat subjective interpretations of available phylogenetic and fossil evidence (as in Benton et al., 2015; Roos et al., 2019; Marjanović, 2021; and most calibrations used by dos Reis et al., 2018). However, dos Reis et al. (2018) based their prior distributions for two nodes - namely the ages of crown Primates and crown Anthropoidea - on the analytical estimates of Wilkinson et al. (2011); nevertheless, we note that the maximum bounds of both of these calibrations (88.6 Ma for crown Primates, 62.1 Ma for crown Anthropoidea) seem implausibly old based on the fossil record (see our proposed calibrations for both of these nodes in “Fossil Calibrations” below). In turn, this may explain why the Late Cretaceous divergence time for crown Primates estimated by dos Reis et al. (2018; 95% Highest Posterior Density [HPD] interval of 70.0-79.2 Ma) is also strongly incongruent with the fossil record, although their estimate for crown Anthropoidea (95% HPD: 41.8-48.3 Ma) is in better agreement with the (very limited) fossil evidence for this node (see “Fossil Calibrations” below).

Here, we take these considerations into account to specify an up-to-date set of well-justified fossil calibrations for divergences within Primates, including several entirely new calibrations. In our initial survey of the literature, we identified numerous recent studies that have formally tested the affinities of various fossil primates and relatives using large morphological and total evidence datasets, many of them using tip-dating (e.g. Dembo et al., 2015, 2016; Herrera and Dávalos, 2016; Gunnell et al., 2018; Ni et al., 2019; Seiffert et al., 2020; Püschel et al., 2021). However, these studies collectively include a relatively limited sample of platyrrhine taxa (the so-called “New World monkeys”). For this reason, we firstly carry out a tip-dating analysis of living and fossil platyrrhines, based on a total evidence dataset that combines 10.2 kilobases (kb) of DNA sequence data from 17 nuclear genes (taken from Woods et al., 2018), and 418 morphological characters (taken from Kay et al., 2019), to identify fossil taxa that can be robustly placed within crown Platyrrhini and that, as a result, can be used to calibrate the ages of specific platyrrhine divergences.

Secondly, we combine our results with other published evidence regarding the fossil record and phylogeny of primates and other mammals to identify calibrations for 27 nodes: four outside Primates (crown Euarchontoglires, crown Glires, crown Euarchonta and crown Primatomorpha), crown Primates itself, and 22 divergences within crown Primates. This is a >50% increase in the number of calibrations compared to other recent broadscale molecular clock analyses of primates (Perelman et al., 2011; Springer et al., 2012; dos Reis et al., 2018; Vanderpool et al., 2020). For each calibration, we follow the best practices recommended by Parham et al. (2012). We provide a minimum bound for each calibration, based on the minimum age of the calibrating specimen, and for 24 of our 27 calibrations we also provide a maximum bound and a suggested prior probability distribution (either uniform or exponential), based on our interpretation of the available phylogenetic evidence and the relative completeness of the known fossil record. To maximize the utility of our calibration list to other researchers, we also identify the age of the oldest member of the sister lineage of the calibrating fossil taxon, as required by the CladeAge method for inferring divergence times (Matschiner et al., 2017; Matschiner, 2019). Finally, we compare our proposed calibrations with those suggested in other recent studies (in particular, Benton et al., 2009, 2015; dos Reis et al., 2018; Roos et al., 2019), highlight cases in which there are major differences, and further justify our proposals.

## 2. Methods

### 2.1 Total evidence tip-dating analyses of Platyrrhini

For the total evidence tip-dating phylogenetic analysis of platyrrhines, we combined existing morphological and molecular data. The morphological dataset is that of Kay et al. (2019), which comprises 418 characters from the dentition, cranium, postcranium and soft tissues, of which 314 are parsimony informative, 34 are autapomorphic, and 70 are invariant; of these 418, 175 represent putative morphoclines that can be ordered. Kay et al. (2019) scored these characters for 16 extant platyrrhine genera, 24 fossil and subfossil primate taxa from South and Central America and the Caribbean, and three extant (*Hylobates*, *Miopithecus* and *Presbytis*) and two fossil (*Aegyptopithecus* and *Catopithecus*) catarrhine genera, plus three fossil stem-anthropoids (*Apidium*, *Proteopithecus* and *Simonsius*) that we specified as outgroups. The molecular dataset comprises DNA sequences from 17 nuclear loci that were selected based on their availability for our extant taxon sample and also for the Jamaican subfossil *Xenothrix* (Perelman et al., 2011; Springer et al., 2012; Woods et al., 2018); these sequences were downloaded from Genbank, aligned using MUSCLE with default settings in MEGA, and concatenated into a single file, resulting in a total of 10244 base pairs of sequence data (Table 1). The morphological and molecular datasets were then merged to produce our initial total evidence dataset.

Some of the fossil taxa present in the total evidence dataset are represented by mandibles and lower dentitions only, whilst for others the maxillary dentition is known but the mandible and lower dentition are currently unknown. Relationships between these two categories of taxa cannot be resolved adequately because they are not scored for any characters in common (Springer et al., 2004). For this reason, we prepared two versions of our total evidence dataset for phylogenetic analysis: a “maxillary” dataset in which the four fossil taxa known only from mandibular specimens (“*Aotus*” *dindensis*, *Mohanamico*, *Nuciruptor* and *Proteropithecia*) were deleted, leaving a total of 44 taxa; and a “mandibular” dataset, in which the five fossil taxa for which the mandible and lower dentition are unknown (*Tremacebus*, *Acrecebus*, *Perupithecus*, *Canaanimico* and *Parvimico*) were deleted, leaving a total of 43 taxa.

Tip-dating analyses of the maxillary and mandibular total evidence datasets were carried out using MrBayes 3.2.7a (Huelsenbeck and Ronquist, 2001; Ronquist et al., 2012b), running on the JASMIN data analysis facility (Lawrence et al., 2013). Given the comparatively large number of autapomorphies (34 out of 348 variable characters), the morphological partition of the total evidence dataset was assigned the Lewis (2001) Mk model with the assumption that variable characters have been scored, i.e. the Mk*v* variant (the 70 constant characters were therefore ignored in the analysis), with an eight category lognormal distribution to model rate heterogeneity between sites (Harrison and Larsson, 2015). For the molecular partition, PartitionFinder 2.2 (Lanfear et al., 2017) was used to identify an appropriate partitioning scheme and substitution model for each partition, with possible models restricted to only those implemented by MrBayes, and with variants that combine a gamma distribution to model rate heterogeneity between sites with a proportion of invariant sites not considered, following the recommendations of Stamatakis (2014). The PartitionFinder analysis used the “greedy algorithm” and assumed linked branch lengths, and the Bayesian Information Criterion (BIC) was used for model selection. The best fitting partitioning scheme and set of substitution models for the molecular partition are shown in Table 1. Because we did not consider models that combine a gamma distribution and a proportion of invariant sites, we increased the number of gamma rate categories from the MrBayes default of four to eight.

**Table 1.**
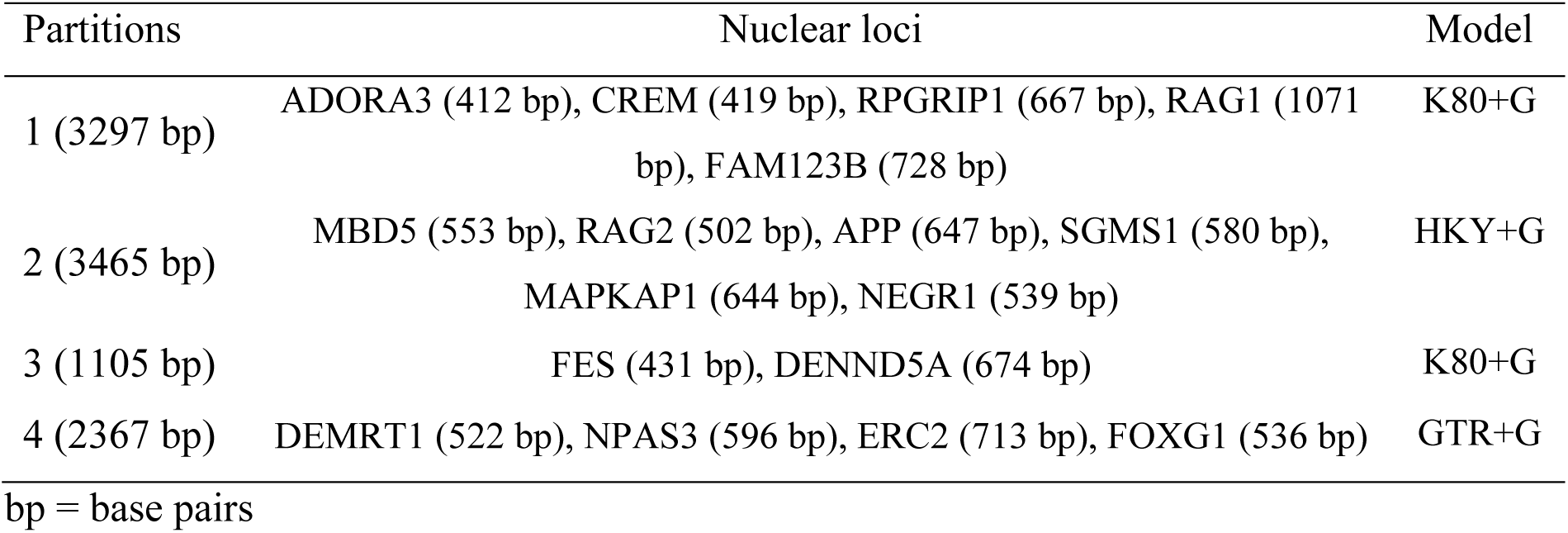
Molecular partitioning scheme for total evidence tip-dating analyses in MrBayes based on PartitionFinder output, Partitions were identified using the Bayesian Information Criterion for model selection and assuming linked branch lengths.

We assigned a fixed age of 0 Ma to all of our extant taxa. For most of our fossil and subfossil taxa, we assigned uniform age ranges corresponding to the entire possible age range of the specimens used for scoring purposes (following Püschel et al., 2020); however, we assigned a fixed age of 33.4 Ma to *Catopithecus* and *Proteopithecus*, all specimens of which are from Quarry L-41 of the Jebel Qatrani Formation in the Fayum Depression of Egypt, which has been assigned an age of 33.4 Ma by Van Couvering and Delson (2020). A full justification of the assigned ages for all of our fossil and subfossil taxa is given in Supplementary Online Material. In an attempt to reduce the time needed to achieve stationarity and convergence between MrBayes runs, we enforced the following topological constraints, corresponding to clades the monophyly of which is uncontroversial: crown Anthropoidea (= all our taxa except the stem anthropoids *Apidium*, *Proteopithecus* and *Simonsius*), crown Catarrhini (= *Hylobates*, *Presbytis* and *Miopithecus*), and crown Cercopithecidae (= *Presbytis* and *Miopithecus*). To assist with rooting, we also specified a partial constraint that enforced monophyly of our extant platyrrhine taxa to the exclusion of our extant non-platyrrhine taxa, but left the position of fossil taxa free to vary. We did not provide node calibrations for any of these clades, i.e., we have used a “tip-dating” approach, rather than “tip-and-node dating” (O’Reilly and Donoghue, 2016). However, following Sallam and Seiffert (2020), we calibrated the age of the root as a truncated normal prior with a minimum of 33.401 Ma (which is only slightly older than the oldest taxa in our dataset, namely *Catopithecus* and *Proteopithecus* - see above), a mean age that is 0.1 Ma older than the age of our oldest fossil taxa (i.e. 33.5 Ma), and a standard deviation of 1.0 Ma.

We applied a single Independent Gamma Rates (IGR) clock model to the entire molecular partition, and a separate IGR model to the morphological partition, with the prior on the clock rate specified using the custom R script of Gunnell et al. (2018), and the prior on the variance left as the MrBayes default. For the tree model, we assumed the Fossilized Birth Death process (Stadler, 2010; Gavryushkina et al., 2014; Heath and Huelsenbeck, 2014), allowing for sampled ancestors, and with diversity sampling assumed (Zhang et al., 2016). Because our focus is Platyrrhini, and our taxa are genus-level (with one possible exception, “*Aotus*” *dindensis*, which nevertheless does not form a clade with extant *Aotus* in several recent phylogenetic analyses; Kay, 2015; Marivaux et al., 2016; Kay et al., 2019; but see Ni et al., 2019), we assumed a sampling probability of 0.727, corresponding to the inclusion of 16 of the 22 extant platyrrhine genera currently recognized. Priors on speciation, extinction and sampling were left as the MrBayes defaults.

MrBayes analyses comprised two independent runs of four MCMC chains (one “cold”, three “heated”, with default heating parameters), run for 40 million generations and sampling trees and other parameters every 5000 generations. Use of Tracer indicated that stationarity and convergence between runs was achieved within four million generations in both analyses; based on this, the first 10% of sampled trees were excluded. The remaining post-burnin trees were summarized in MrBayes using majority rule consensus (following the recommendations of O’Reilly and Donoghue, 2018), rather than a Maximum Clade Credibility tree, with clades retained if they were present in less than 50% of the post-burnin trees provided they were compatible with the 50% majority rule consensus topology (using the “contype=allcompat” command in MrBayes). Support values were calculated as Bayesian posterior probabilities (BPPs). Selected estimates of divergence times in the maxillary and mandibular analyses are given in Table 2.

### 2.2 Identification of fossil calibrations

We used the results from our total evidence tip-dating analyses of platyrrhines, in combination with the published literature, to identify a set of robust fossil calibrations within crown Platyrrhini. Based on published studies, we also identified fossil calibrations for other divergences within crown Primates, for crown Primates itself, and for four other divergences within Euarchontoglires: crown Euarchontoglires, crown Glires, crown Euarchonta, and crown Primatomorpha. In identifying appropriate fossil calibrations, wherever possible we based our decisions on the results of formal, algorithmic phylogenetic analyses that have robustly tested the affinities of relevant fossil taxa, using the following hierarchy (from what we consider to be the most robust analyses to the least robust analyses): total evidence tip-dating analyses; undated total evidence analyses; undated analyses with a molecular scaffold; morphology-only tip-dating analyses; undated morphology-only analyses. In a few cases, we have proposed calibrations that are not based on evidence presented in formal phylogenetic analyses; for most of these, we have based our decisions on the presence of one or more morphological synapomorphies that clearly support assignment of that fossil taxon to a particular clade.

In general, we have followed the recommendations of Parham et al. (2012) for “best practices” for fossil calibrations; these include identifying a specific fossil specimen for each calibration, providing a full phylogenetic justification for using a particular fossil taxon (following our general approach listed above), discussing (where relevant) differences between morphological and molecular phylogenetic analyses, giving detailed locality and stratigraphic information for the calibrating specimen, and explaining clearly how this translates to a particular fixed age or age range. For each calibration, we provide a minimum bound, which we argue can reasonably be viewed as “hard” (i.e. zero probability of the divergence being younger than this). For most calibrations, we also propose a maximum bound, which should be viewed as “soft” (i.e., with a small probability that the divergence is older than this). Where we have provided a minimum and a maximum bound, we also propose a prior probability distribution. For most calibrations, we propose a uniform distribution, in which all ages between the minimum and the maximum bound have equal probability, which we consider to be most appropriate in cases where the fossil record is obviously highly incomplete (Ho and Phillips, 2009). For a few calibrations, however, we consider that the fossil record is sufficiently complete to be relatively confident the true age of divergence is close to the minimum bound; in such cases, we suggest that this should be modelled as an offset exponential distribution, with the minimum bound as the offset, and the shape of the exponential distribution specified such that there is a 5% probability of the divergence being older than the maximum bound (Ho and Phillips, 2009). In each case, we explain in detail why we consider a uniform or an offset exponential prior distribution is appropriate.

Our list of a minimum and (where relevant) a maximum bound for a calibrated node reflects standard practice in node-dating analyses. However, Matschiner et al. (2017) proposed the “CladeAge” method (see also Matschiner, 2019), in which information about the oldest fossils for particular clades is combined with estimates of sampling and diversification rates to construct prior distributions on the ages of those clades. The simulation study of Matschiner (2019) suggests that the CladeAge method is more robust to model violations than are standard node dating-analyses that use the Fossilized Birth Death (FBD) model (Stadler, 2010; Gavryushkina et al., 2014; Heath and Huelsenbeck, 2014). The CladeAge method requires information on the oldest member of each clade present in a phylogeny, not just of the oldest of the daughter clades descending from a particular node (as required in “standard” node-dating). Therefore, to maximize the utility of our calibration list to other researchers, and to allow use of the CladeAge method, for each calibrated node, we provide ages for the oldest member of each daughter lineage descending from that node. It should be noted the CladeAge method also requires estimates of sampling and diversification rates, which we do not give here (such rates can be estimated using programs such PyRate; Silvestro et al., 2014, 2019a). Table 3 summarizes our full calibration list, including minimum and (where relevant) maximum bounds, suggested prior age distributions, and calibrations in a format suitable for use in the CladeAge method.

## 3. Results

Relationships among extant platyrrhines recovered by tip-dating analysis of the maxillary and mandibular total evidence datasets, using the Fossilized Birth Death (FBD) model, are highly congruent (Figures 1-2). Both analyses place all pre-Laventan (i.e., older than ∼13.8-11.8 Ma; Prevosti and Forasiepi, 2018) fossil primates from Central and South America outside crown Platyrrhini, with one potential exception: *Proteropithecia* from the Collón Curá Formation of Argentina (present in the mandibular dataset only), which is very weakly supported (BPP = 0.30) as a stem pitheciine. The age of *Proteropithecia* is poorly constrained (19.76-10.4 Ma old; see Supplementary Online Material), but may still predate the Laventan. However, examination of the individual post-burnin trees from the mandibular analysis reveals slightly stronger support (BPP = 0.34) for placing *Proteropithecia* outside crown Platyrrhini (note that this is not seen in Figure 1 because the post-burnin trees that find this relationship conflict with the overall 50% majority rule consensus topology).

**Figure 1.**
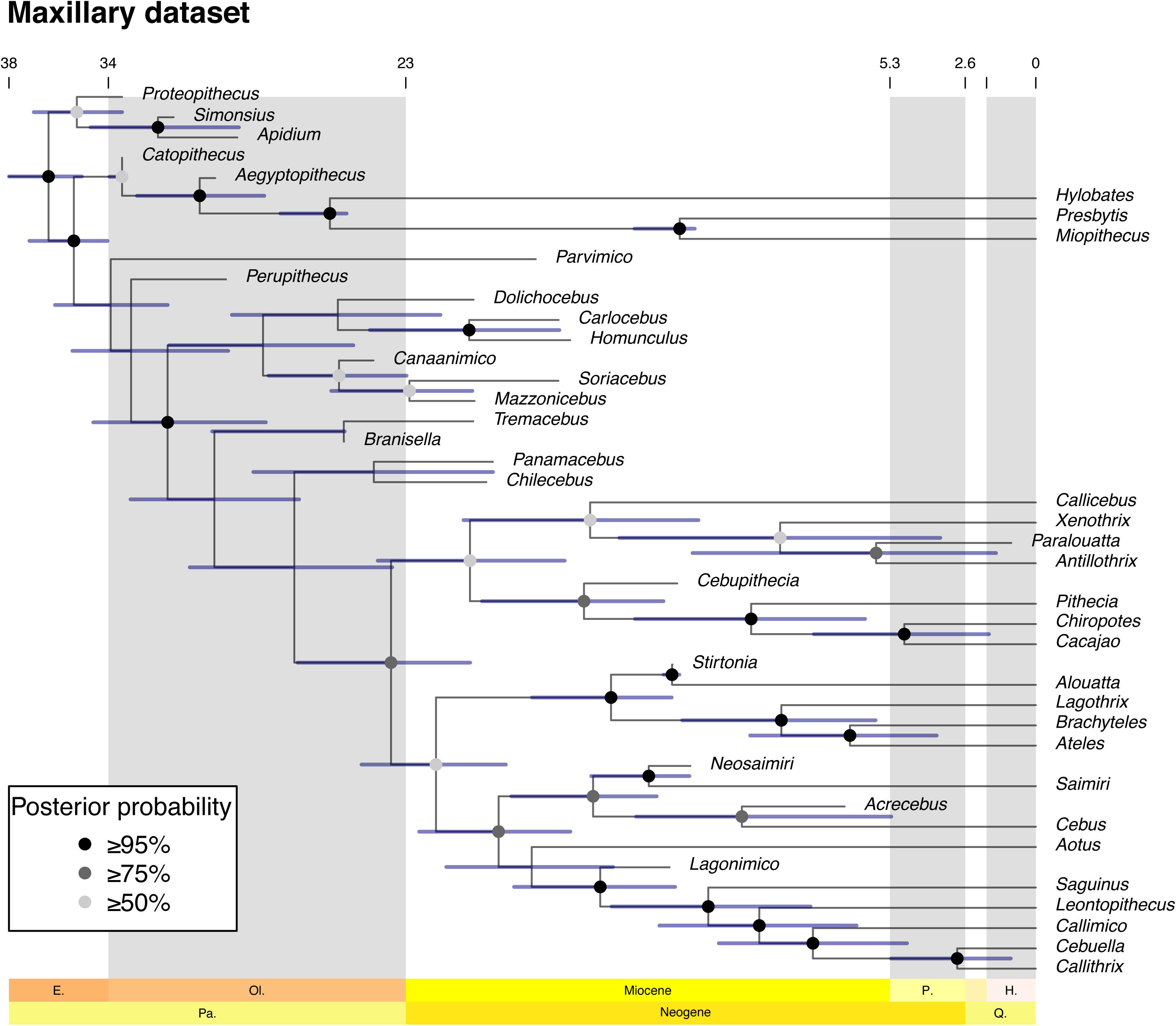
“Allcompat” majority rule consensus of post-burn-in trees that results from tip-dating Bayesian analysis of our “maxillary” total evidence dataset using MrBayes. Divergence dates are median posterior estimates, and blue bars are 95% Highest Posterior Density (HPD) intervals.

**Figure 2.**
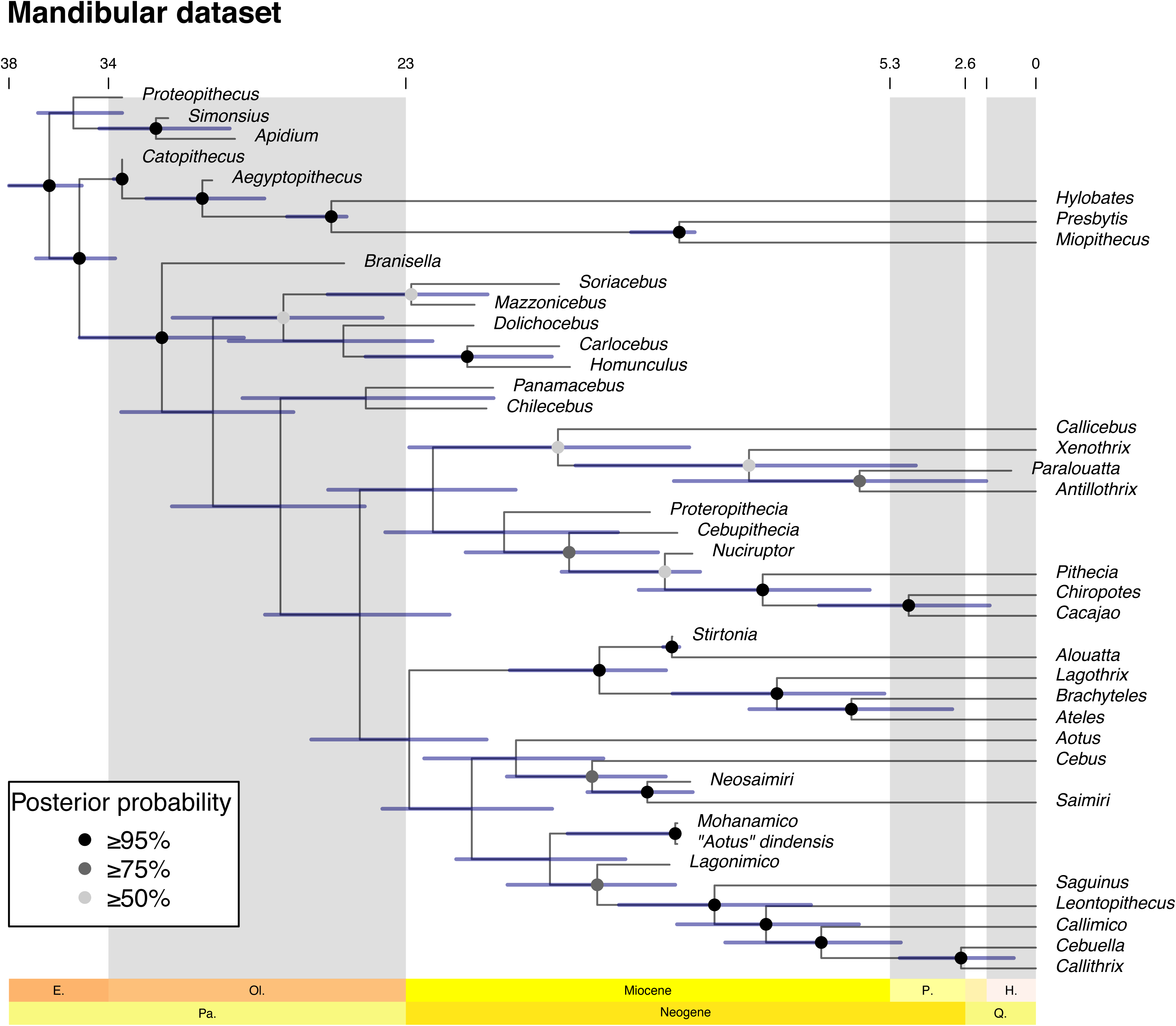
“Allcompat” majority rule consensus of post-burn-in trees that results from tip-dating Bayesian analysis of our “mandibular” total evidence dataset using MrBayes. Divergence dates are median posterior estimates, and blue bars are 95% Highest Posterior Density (HPD) intervals.

Of particular note is the fact that *Panamacebus*, which was identified as a cebid in its original description (Bloch et al., 2016), is recovered outside the crown clade in both the mandibular and maxillary analyses, where it forms a very weakly supported clade with *Chilecebus* (BPP of 0.38 in the mandibular analysis, and 0.41 in the maxillary analysis). Monophyly of crown Platyrrhini excluding *Panamacebus* is strongly supported in the maxillary analysis (BPP = 0.82), although not in the mandibular analysis (BPP = 0.36), likely because of the unstable position of *Proteropithecia* discussed above. Among the other fossil stem platyrrhines, there is strong support for *Homunculus*+*Carlocebus* (BPP of 0.99 in both analyses), and somewhat weaker support for *Mazzonicebus*+*Soriacebus* (BPP of 0.69 in the mandibular analysis, and 0.70 in the mandibular analysis), but most other relationships otherwise received very weak support (BPP <0.5). Among these very weakly supported relationships is a position for the early Miocene (19.6-16.4 Ma) *Parvimico* as sister to all other crown and stem platyrrhines, with the early Oligocene (29.68-29.52 Ma) *Perupithecus* the next taxon to branch (these two taxa are present in the maxillary analysis only).

Within crown Platyrrhini, the topology in both analyses is congruent with recent molecular phylogenies in supporting a basal split between Pitheciidae (here also including the three Caribbean subfossil taxa *Xenothrix*, *Antillothrix*, and *Paralouatta*) and a clade comprising the remaining extant families (Atelidae, Aotidae, Callitrichidae and Cebidae); these two clades receive weak-to-moderate support in the maxillary analysis (BPP of 0.60 and 0.62, respectively) but not in the mandibular analysis (BPP of 0.32 and 0.36, respectively), again likely due to instability in the position of *Proteropithecia* in the latter analysis. Both analyses also agree in placing Atelidae sister to a clade comprising Aotidae (*Aotus*), Callitrichidae and Cebidae, although again support for the latter clade is stronger in the maxillary analysis (BPP = 0.79) than the mandibular analysis (BPP = 0.49). The analyses disagree on the position of Aotidae (*Aotus*), which is sister to Cebidae in the mandibular analysis (BPP = 0.34) but sister to Callitrichidae in the maxillary analysis (BPP = 0.46), but support is very weak in both cases.

Turning now to the Laventan fossil taxa, there is strong support for placing *Stirtonia* as sister to *Alouatta* within crown Atelidae (BPP = 1.00 in both analyses), *Neosaimiri* as sister to *Saimiri* within crown Cebidae (BPP = 1.00 in both analyses), and *Lagonimico* as a stem callitrichid (BPP of 1.00 in the maxillary analysis, and 0.95 in the mandibular analysis). In the mandibular analysis, there is also strong support for *Cebupithecia* and *Nuciruptor* forming a clade with extant pitheciines (BPP = 0.92), whereas there is somewhat weaker support for placing *Cebupithecia* with extant pitheciines in the maxillary analysis (BPP = 0.77); *Nuciruptor* is known from lower jaw material only, and so was not included in the latter analysis. The other Laventan taxa known solely from lower jaw material, namely “*Aotus*” *dindensis* and *Mohanamico*, form a strongly supported (BPP = 1.00) clade in the mandibular analysis, which is in turn part of a weakly supported (BPP = 0.36) clade that also includes Aotidae (*Aotus*), Callitrichidae and Cebidae. The Huayquerian (9.0-5.28 Ma) *Acrecebus*, which is known from a single upper molar, receives fairly strong support (BPP = 0.87) for a position as sister to *Cebus* in the maxillary analysis.

Strikingly, both the maxillary and mandibular analyses recover a clade comprising the Caribbean subfossil *Xenothrix*, *Antillothrix*, and *Paralouatta*, although support for this is relatively weak (BPP of 0.59 in the maxillary analysis, and 0.58 in the mandibular analysis). Both analyses also place this Caribbean clade sister to *Callicebus*, but support for this relationship is again quite weak (BPP of 0.58 in the maxillary analysis, and 0.50 in the mandibular analysis).

Focusing now on divergence times (see Table 2), the oldest split within total-clade Platyrrhini found here is between *Parvimico* (present in the maxillary analysis only) and the remaining taxa, with a median estimated age of 33.8 Ma and 95% HPD of 31.7-35.8 Ma. The maxillary and mandibular analyses both place the origin of crown Platyrrhini in the late Oligocene or early Miocene, with a median estimate of 23.6 Ma and 95% HPD of 20.7-27.0 Ma in the maxillary analysis, and a median estimate of 24.7 Ma and 95% HPD of 21.4-28.2 Ma in the mandibular analysis. Among the fossil taxa, the very short temporal branch (0.01 Ma) leading to *Stirtonia* in both analyses is congruent with its being ancestral to the extant *Alouatta*, but there are no other plausible ancestor-descendant relationships among our fossil taxa. The divergence between *Callicebus* and the Caribbean clade is dated to the early-to-middle Miocene in both analyses (median estimate of 16.3 Ma and 95% HPD of 12.3-20.9 Ma in the maxillary analysis; median estimate of 17.5 Ma and 95% HPD of 12.7-22.9 Ma in the mandibular analysis). The first divergence within the Caribbean clade itself (the split between *Xenothrix* and *Antillothrix*+*Paralouatta*) has important biogeographical implications, but is only weakly constrained to some point from the middle Miocene to the Pliocene here, with a median estimate of 9.3 Ma and 95% HPD of 3.5-15.2 Ma in the maxillary analysis, and a median estimate of 10.5 Ma and 95% HPD of 4.4-16.8 Ma in the mandibular analysis. Selected estimated divergence times are summarized in Table 2.

**Table 2.** Divergence dates estimated for selected clades in our total evidence tip-dating analyses. The “maxillary” analysis excluded fossil taxa known only from mandibular specimens; the “mandibular” analysis excluded fossil taxa for which the mandible and lower dentition are unknown. Values are median posterior estimates, followed by 95% Highest Posterior Density (HPD) intervals in brackets.

| Clade | Split | Maxillary | Mandibular |
| --- | --- | --- | --- |
| crown Platyrrhini | Pitheciidae-<br>(Aotidae+Atelidae+Callitrichidae+Cebidae) | 23.6 (20.7-27.0) | 24.7 (21.4-28.2) |
| crown Pitheciidae | Pitheciinae-Callicebinae | 20.7 (17.2-24.1) | 22.0 (19.0-25.9) |
| crown Pitheciinae | *Pithecia*-(*Chiropotes*+*Cacajao*) | 10.4 (6.2-14.7) | 10.0 (6.1-14.5) |
| *Callicebus* + Caribbean<br>platyrrhines | *Callicebus*-(*Xenothrix*+*Antillothrix*+*Paralouatta*) | 16.3 (12.3-20.9) | 17.5 (12.7-22.9) |
| Caribbean platyrrhines | *Xenothrix*-(*Antillothrix*+*Paralouatta*) | 9.3 (3.5-15.2) | 10.5 (4.4-16.8) |
| Atelidae + Aotidae +<br>Callitrichidae + Cebidae | Atelidae-(Aotidae+Callitrichidae+Cebidae) | 21.9 (19.4-24.6) | 22.9 (20.1-26.5) |
| crown Atelidae | Alouattinae-Atelinae | 15.5 (13.3-18.4) | 16.0 (13.5-19.2) |
| crown Atelinae | *Lagothrix*-(*Brachyteles*+*Ateles*) | 9.3 (5.9-12.9) | 9.5 (5.5-13.3) |
| crown Callitrichidae | *Saguinus-(Leontopithecus+Cebuella+Callithrix+Callimico)* | 12.0 (8.2-15.5) | 11.7 (8.2-15.2) |
| crown Cebidae | Cebinae-Saimirinae | 16.2 (13.9-19.2) | 16.2 (13.5-19.3) |

## 4. Discussion

To date, there have been relatively few published total evidence phylogenetic analyses focused on platyrrhine relationships. Several recent studies (Kay, 2015; Bloch et al., 2016; Marivaux et al., 2016; Kay et al., 2019) have used an alternative approach for incorporating molecular information, namely constraining relationships among modern taxa to match current molecular phylogenies (a molecular “scaffold”), but this prevents the morphological and molecular data from interacting synergistically (Hermsen and Hendricks, 2008).

Schrago et al. (2013) did present a total evidence analysis of Platyrrhini, and used a novel approach to infer divergence dates by first carrying out an undated total evidence analysis, and then performing a relaxed clock analysis of morphological data only on the total evidence topology, with age estimates from a separate molecular clock analysis used as node age priors. However, this approach lacks a key aspect of tip-dating, namely that the stratigraphic ages of the fossil terminals can influence their phylogenetic position (Bapst et al., 2016; Turner et al., 2017; Lee and Yates, 2018; Beck and Taglioretti, 2020; King and Beck, 2020; Beck et al., 2021; King, 2021). The morphological clock analysis of Schrago et al. (2013) also assumed a Yule prior on tree shape, which has been shown to lead, in at least some cases, to unrealistically ancient divergence dates (Condamine et al., 2015), although we note that the dates calculated by Schrago et al. (2013) in this analysis are in fact largely congruent with those presented here (Table 2). The sampling of fossil platyrrhines by Schrago et al. (2013) was also relatively restricted, and lacked a number of taxa that have been consistently associated with crown lineages, such as the *Stirtonia*, *Cebupithecia*, *Neosaimiri*, and *Lagonimico* (Kay, 2015). Indeed, only one fossil taxon (*Proteropithecia*) fell within crown Platyrrhini (as a stem pitheciine) in the analysis of Schrago et al. (2013), and so the results of this analysis are of limited use in identifying robust fossil calibrations for divergences among crown platyrrhines. In addition, our own results raise questions about whether *Proteropithecia* is indeed a pitheciine (discussed in more detail below).

Finally, Silvestro et al. (2019b) carried out a tip-dating analysis of platyrrhines that included 34 fossil taxa, using the FBD model, but the fossil taxa were not represented by character data (living taxa were represented by combined nuclear and mitochondrial sequence data only), and their relationships were constrained a priori; thus, their phylogenetic position was not free to vary, and could not be informed by morphological data. Perhaps most significantly, Silvestro et al. (2019b) assumed the “Long Lineage” hypothesis, which considers that pre-Laventan platyrrhines from Patagonia are members of the crown clade (Rosenberger et al., 2009; Rosenberger, 2010, 2011, 2019; Rosenberger and Tejedor, 2013), when constraining the relationships of their fossil taxa; the “Long Lineage” hypothesis is not supported by our analyses (see below).

Published total evidence tip-dating analyses of broad primate phylogeny, meanwhile, have typically included a relatively limited sampling of platyrrhines (e.g. Gunnell et al., 2018; Seiffert et al., 2020). An exception is that of Ni et al. (2019: fig. S1), which included a relatively dense sampling of fossil platyrrhines, as well as other primates, and which used a similar approach to that here (specifically, the Fossilized Birth Death tree model in combination with the IGR clock model, as implemented in MrBayes), but with a different dataset (1186 morphological characters from the dentition, cranium, postcranium and soft tissues; 658 SINE and LINE insertion characters) that was focused on relationships within Haplorhini as a whole. However, of the 11 fossil platyrrhines included by Ni et al. (2019 fig. S1), only two fell within crown Platyrrhini in their analysis: *“Aotus” dindensis* as sister to the extant *Aotus azarae*, and *Mohanamico hershkovitzi* as a stem callitrichid. Several other recent analyses (Kay, 2015; Bloch et al., 2016; Marivaux et al., 2016; Kay et al., 2019) have failed to support “*A.*” *dindensis* as an aotid, and have also placed two fossil taxa that Ni et al. (2019 fig. S1) found to be stem platyrrhines - namely *Stirtonia* and *Neosaimiri* - within crown Platyrrhini. Like Schrago et al. (2013), Ni et al. (2019 fig. S1) also did not include several other fossil taxa that have been suggested to fall within crown Platyrrhini, namely *Proteropithecia*, *Cebupithecia*, *Nuciruptor* and *Lagonimico* (Kay, 2015).

The total evidence tip-dating analyses we present here should therefore be of interest because they are the first to include a diverse sampling of fossil platyrrhines, including putative members of many crown lineages. As such, they should help identify robust calibrations within crown Platyrrhini for use in molecular clock analyses, as well as providing their own estimates of platyrrhine relationships and divergence times.

Relationships among the extant platyrrhines recovered in our both our total evidence tip-dating analyses are strongly congruent with most recent molecular analyses of Platyrrhini in supporting a basal split between Pitheciidae and the remaining families, with Atelidae sister to a Aotidae+Callitrichidae+Cebidae clade (Perelman et al., 2011; Perez et al., 2012; Springer et al., 2012; dos Reis et al., 2018; Valencia et al., 2018; Woods et al., 2018; but see X. Wang et al., 2019). Our two analyses differ over the precise position of Aotidae (= *Aotus*), with the mandibular analysis placing this family sister to Cebidae but the maxillary instead placing it sister to Callitrichidae, but with very weak support (BPP <0.5) in both cases. Failure of both of our analyses to unambiguously resolve the precise branching pattern between Aotidae, Callitrichidae and Cebidae is perhaps unsurprising as much larger molecular sequence datasets also fail to do this (Osterholz et al., 2009; Perez et al., 2012; Valencia et al., 2018; Schrago and Seuánez, 2019; X. Wang et al., 2019; Vanderpool et al., 2020). Relationships among extant members of Callitrichidae, Atelidae and Pitheciidae are congruent with recent molecular analyses (Perelman et al., 2011; Springer et al., 2012; Buckner et al., 2015; dos Reis et al., 2018; Garbino and Martins-Junior, 2018).

In terms of relationships among our non-living taxa, perhaps the most significant aspect of both analyses is the fact that all pre-Laventan (middle Miocene) South American primate taxa (all of which come from Patagonia in southern South America) included in our analyses fall outside crown Platyrrhini, with the possible exception of *Proteropithecia*. Of particular interest is the position of the early Miocene (21.1-18.748 Ma) *Panamacebus*, which was originally described as a cebid by Bloch et al. (2016), and which has been recovered within Cebidae in several published phylogenetic analyses that have used a molecular scaffold approach (Bloch et al., 2016; Marivaux et al., 2016; Kay et al., 2019). Here, both of our total evidence tip-dating analyses instead place *Panamacebus* outside crown Platyrrhini, in a very weakly supported (BPP <0.5) clade with *Chilecebus*. Thus, our analyses do not support the “Long Lineage” hypothesis (Rosenberger et al., 2009; Rosenberger, 2010, 2011, 2019; Rosenberger and Tejedor, 2013), which posits that the pre-Laventan taxa are early members of crown platyrrhine lineages.

The case of *Proteropithecia* from the Collón Curá Formation of Argentina warrants further discussion. Firstly, the age of *Proteropithecia* is poorly constrained, to between 19.76 and 10.4 Ma (see Supplementary Online Material), and so it may in fact postdate the Laventan (which is ∼13.8-11.8 Ma; Prevosti and Forasiepi, 2018). Secondly, although the majority rule consensus from our mandibular analysis placed *Proteropithecia* within crown Platyrrhini, as a stem pitheciine (congruent with current hypotheses), support for this relationship was very weak (BPP = 0.30), and in fact 34% of the post-burnin trees (equivalent to a BPP of 0.34) placed *Proteropithecia* outside crown Platyrrhini. To our knowledge, the pitheciine status of *Proteropithecia* has been uncontroversial (Kay et al., 2013; Rosenberger and Tejedor, 2013; Kay, 2015; Tejedor and Novo, 2016; Rosenberger, 2020), although Kay et al. (1998) noted some notable dental differences between *Proteropithecia* and living pitheciines in their original description of this taxon. We do not attempt a detailed reassessment of the affinities of *Proteropithecia*, but merely note that our results raise the possibility that this taxon may be a stem platyrrhine convergent on pitheciines (as has also been proposed for *Soriacebus*; Kay, 1990, 2010, 2015; Kay et al., 2013). If so, this would remove the last direct biogeographical link between the Patagonian platyrrhines (all of which would therefore seem to be stem taxa) and crown Platyrrhini, the known fossil record of which is confined to northern South America.

Another notable aspect of both of our analyses is recovery of a clade (sister to *Callicebus*) comprising the three Caribbean subfossil taxa *Xenothrix*, *Antillothrix* and *Paralouatta*. This clade was also found by Marivaux et al. (2016), but not by Kay (2015) or Kay et al. (2019), even though we used the latter’s morphological matrix here; however, all of these previous studies were based on parsimony analysis of a morphological matrix with a molecular scaffold enforced, which precludes synergistic interactions between morphological and molecular data and which does not take into account temporal information, unlike the approach used here. The Caribbean clade receives weak support in our analyses (BPP of 0.50 in the mandibular analysis, and 0.58 in the maxillary analysis), but it is of interest because it implies that the presence of these genera in the Caribbean may be the result of a single dispersal event, as also recently suggested for Caribbean caviomorph rodents based on DNA evidence (Woods et al., 2021; although the presence of oryzomyin muroids and an apparent geomorph still indicates multiple dispersals by rodents from mainland South America to Caribbean landmasses; Marivaux et al., 2021). Of the three Caribbean taxa, DNA sequence data is currently only available for *Xenothrix* (Woods et al., 2018); obtaining molecular data (e.g., ancient DNA or protein sequences) from *Antillothrix*, *Paralouatta*, and also *Insulacebus* (Cooke et al., 2011), will allow rigorous testing of this hypothesis.

Based on our results, the following four South American fossil primate taxa can be identified as representing robust calibrations for informing the minimum bound on divergences within Platyrrhini: *Stirtonia*, which is strongly supported as a stem alouattine, and so provides a minimum bound on age of the Alouattinae-Atelinae split; *Neosaimiri*, which is strongly supported as a stem saimirine, and so provides a minimum bound on the age of the Cebinae-Saimirinae split; *Cebupithecia*, which is strongly supported as a pitheciine, and so provides a minimum bound on the age of Pitheciinae-Callicebinae split; and *Lagonimico*, which is strongly supported as a stem callitrichid, and so provides a minimum bound on the divergence of Callitrichidae from its sister family (which varies depending on the precise relationship of Aotidae to Callitrichidae and Cebidae). Each of these calibrations is discussed in more detail below (see “Fossil Calibrations”). We note here, however, that the tip-dating total evidence analysis of Ni et al. (2019: fig. S1) suggests a very different set of relationships, with *Stirtonia* and *Neosaimiri* both placed outside crown Platyrrhini (*Cebupithecia* and *Lagonimico* were not included).

Studies suggest that accurate estimates of divergence times under the Fossilized Birth Death model require dense sampling of fossil taxa (Klopfstein et al., 2019; O’Reilly and Donoghue, 2019). The morphological dataset we used here, namely that of Kay et al. (2019), includes most named South American primate genera (note that *Szalatavus*, *Killikaike*, and *Laventiana* have been argued to be synonyms of *Branisella*, *Homunculus*, and *Neosaimiri* respectively; Kay, 2015), but this is a reflection of the overall paucity of the known fossil record (although this continues to improve thanks to ongoing, extensive efforts by a number of different research teams; Kay et al., 2019; Marivaux et al., 2020a; Seiffert et al., 2020; Antoine et al., 2021; Novo et al., 2021). However, this dataset includes a limited selection of fossil catarrhines and stem anthropoids, some of which (e.g., *Talahpithecus*, *Proteopithecus*, and oligopithecids) may be of particular relevance for understanding the origin of platyrrhines, including the timing of their arrival in South America (Bond et al., 2015). For this reason, the divergence times estimated in our total evidence tip-dating analyses should be treated with caution. Nevertheless, our maxillary analysis suggests that platyrrhines arrived in South America between 36.8 and 31.7 Ma, based on the maximum estimate for the Platyrrhini-Catarrhini split (95% HPD: 36.8-33.9 Ma) and the minimum estimate for the divergence of the earliest diverging South American taxon (*Parvimico*, known only from a single upper molar, and hence not present in the mandibular analysis) in our dataset (95% HPD: 35.8-31.7 Ma). Strikingly, this is very similar to Seiffert et al.’s (2020) estimate for the timing of dispersal of a second South American primate taxon, the parapithecid *Ucayalipithecus*, namely 35.1-31.7 Ma, which these authors noted coincides with a major drop in global sea levels (Miller et al., 2008).

Our late Oligocene-early Miocene estimate for the age of crown Platyrrhini is similar to estimates from several previous molecular clock analyses (Perelman et al., 2011; Springer et al., 2012; dos Reis et al., 2018; Woods et al., 2018; X. Wang et al., 2019), as well as from the novel two stage molecular and morphological clock analysis of Schrago et al. (2013), and from the total evidence tip-dating analysis of Ni et al. (2019). It is worth noting that this is similar to the crown age of another major clade of predominantly South and Central American mammals, namely the marsupial family Didelphidae (opossums), as estimated by some molecular clock (node-dating) analyses (Jansa et al., 2014) - although others suggest a somewhat earlier age (Steiner et al., 2005; Meredith et al., 2011; Mitchell et al., 2014; Vilela et al., 2015) -, and also by total evidence clock (combined tip-and-node dating; O’Reilly and Donoghue, 2016) analyses (Beck and Taglioretti, 2020; Beck et al., 2021).

Node-dated molecular clock estimates for the age of crown Caviomorpha are markedly older, typically mid-to-late Eocene or earliest Oligocene (Sallam et al., 2009; Rowe et al., 2010: table 1; Meredith et al., 2011; Upham and Patterson, 2012, 2015; Voloch et al., 2013; Álvarez et al., 2017; Woods et al., 2021). Several of these studies (Voloch et al., 2013; Upham and Patterson, 2015; Álvarez et al., 2017) have calibrated the deepest divergences within crown Caviomorpha based on fossil taxa from Contamana in Peru, which have been dated to ∼41 Ma and which have been referred to crown clades by some authors (Antoine et al., 2012; Boivin et al., 2017). However, this reported age has been questioned (Campbell et al., 2021), and, in any case, the recent phylogenetic study of Boivin et al. (2019) suggests that the Contamana rodents fall outside the crown clade. However, definitive crown caviomorphs have been reported from the Tinguiririca fauna of Chile (Bertrand et al., 2012), and appear to be 37.5-31.5 Ma based on radiometric dating (Wyss et al., 1990, 1993; Flynn et al., 2003; Bertrand et al., 2012), indicating that crown Caviomorpha had begun to diversify by the earliest Oligocene at the latest. Campbell et al. (2021: fig. S7) presented a morphological tip-dating analysis of Caviomorpha (based on the matrix of Marivaux et al., 2020b) in which the ages of fossil taxa from localities of controversial age (at Santa Rosa, Shapaja, and Contamana; Campbell et al., 2021) were allowed to vary between 56 and 0 Ma. In this analysis, the median estimate for the age of crown Caviomorpha was 40.2 Ma, with a 95% HPD of 43.5-37.5 Ma. Collectively, then, current evidence suggests that crown Caviomorpha is at least 10 million years older than crown Platyrrhini.

The tip-dating analysis of Campbell et al. (2021: fig. S7) suggests that the dispersal of Caviomorpha to South America occurred after 44.5 Ma but before 37.5 Ma. This is close to, but does not overlap with, our composite estimate for the timing of dispersal of platyrrhines (36.8-31.7 Ma), or Seiffert et al.’s (2020) estimate for the dispersal of the parapithecid lineage represented by *Ucayalipithecus* (35.1-31.7 Ma). However, the accuracy of divergence dates estimated by tip-dating approaches needs further exploration (Beck and Lee, 2014; Drummond and Stadler, 2016; Puttick et al., 2016; Ronquist et al., 2016; Parins-Fukuchi and Brown, 2017; Luo et al., 2019; King and Beck, 2020; Püschel et al., 2020; Simões et al., 2020), and molecular clock studies suggest that the dispersals of platyrrhines and caviomorphs could have been coincident in time (Poux et al., 2006; Loss-Oliveira et al., 2012).

As noted above, monophyly of the Caribbean genera *Xenothrix*, *Antillothrix* and *Paralouatta* raises the possibility that they may be the result of a single dispersal event from the South American mainland; if so, our divergence estimates permit a wide range of ages for this event, spanning from the earliest Miocene to the late Pliocene (22.9-3.5 Ma). Although poorly constrained, this nevertheless overlaps with the inferred timing of dispersal of caviomorph rodents to Caribbean landmasses based on the molecular clock (node-dating) analysis of Woods et al. (2021), which is 21.7-7.1 Ma. These estimates are therefore permissive of synchronous dispersals by platyrrhines and caviomorphs from mainland South America to Caribbean landmasses.

## 5. Fossil calibrations

**Table 3.**
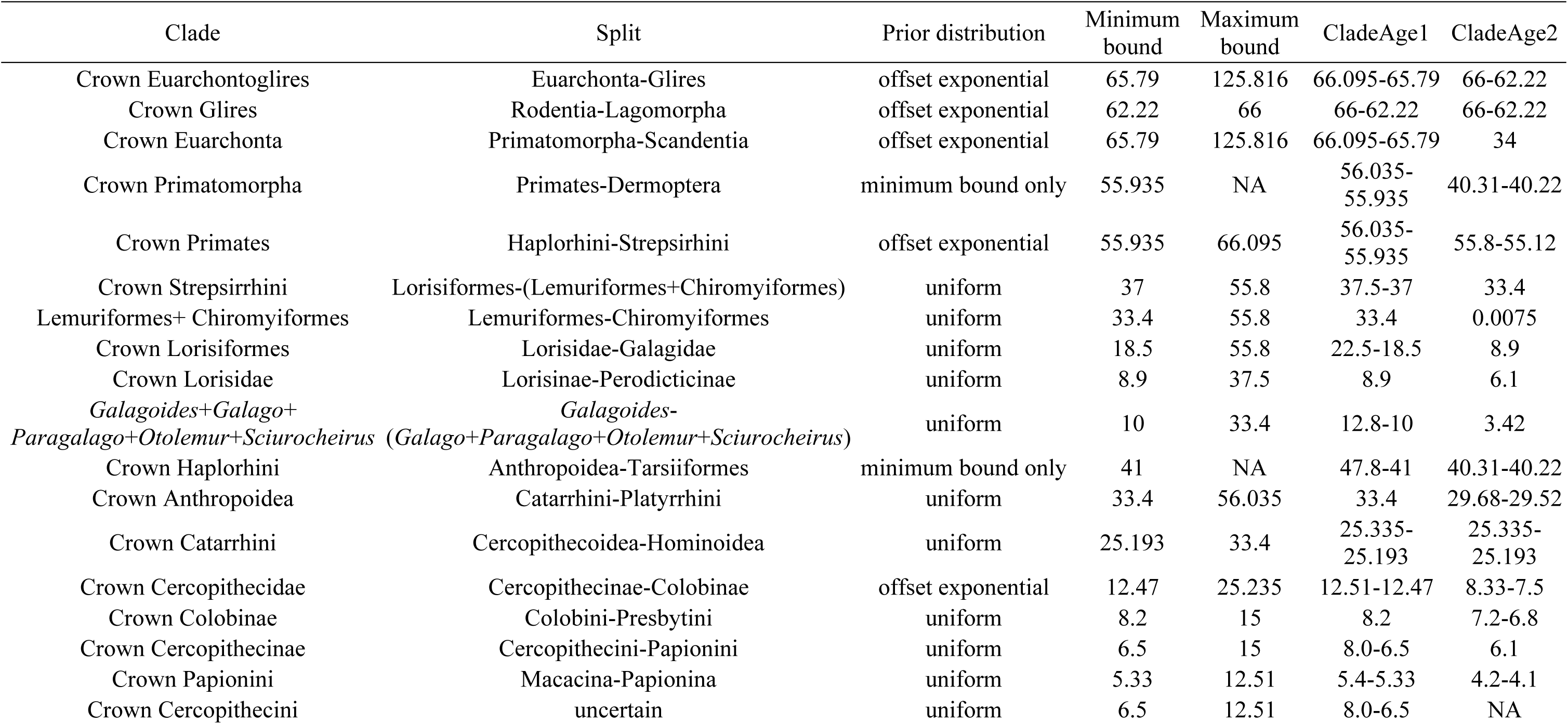
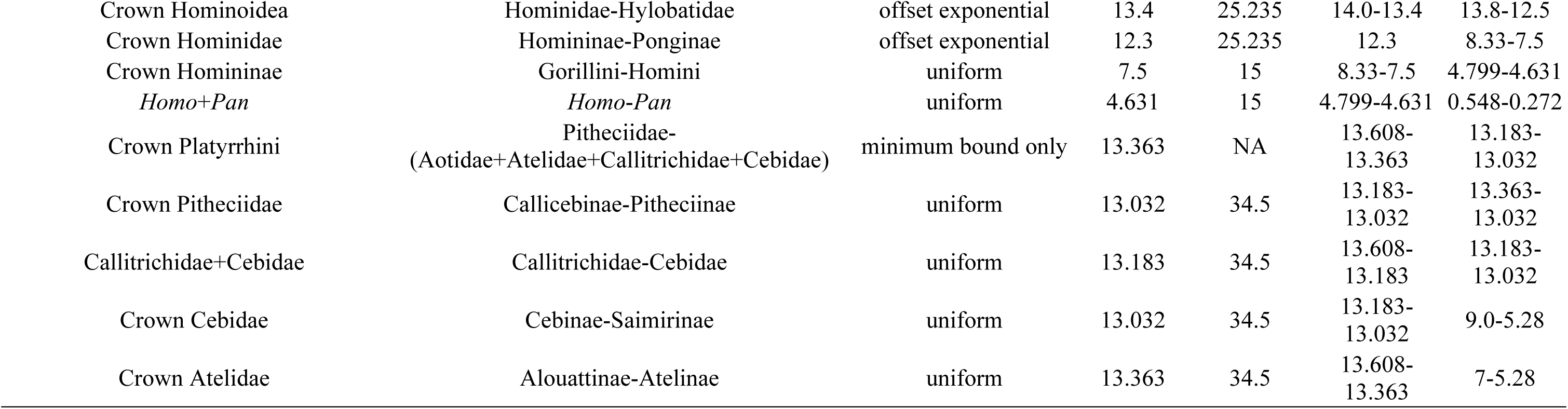
Summary of fossil calibrations proposed here. Values are in millions of years. “CladeAge1” and “CladeAge2” are the ages of the oldest representatives of the two daughter lineages originating after a particular split. See text for full details.

### 5.1 Crown Euarchontoglires = Euarchonta-Glires split

<u>Calibrating taxon</u> Purgatorius mckeeveri

<u>Specimen</u> UCMP 150021, an isolated left m2, from Harley’s Point locality in the lowermost part of the Tullock Member of the Fort Union Formation in Montana, USA (Wilson Mantilla et al., 2021).

<u>Phylogenetic justification</u> *Purgatorius mckeeveri* is the oldest known member of “Plesiadapiformes”, a likely non-monophyletic grade of fossil euarchontans (Silcox et al., 2017). Some phylogenetic analyses focused on deep relationships within Eutheria have recovered *Purgatorius* outside Placentalia (e.g., Wible et al., 2007, 2009; Goswami et al., 2011). However, recent phylogenetic analyses specifically intended to resolve euarchontan relationships consistently place *Purgatorius* and other “plesiadapiforms” within crown Euarchonta, and specifically closer to Primates and Dermoptera than to Scandentia, although the precise affinities of “plesiadapiforms” vary in these analyses (e.g., Bloch et al., 2007; Chester et al., 2015, 2017; Li and Ni, 2016; Ni et al., 2016; Silcox et al., 2017; Gunnell et al., 2018; Morse et al., 2019; Seiffert et al., 2020): different “plesiadapiform” taxa may represent stem-members of Primates and/or Dermoptera, or they may fall outside crown Primatomorpha (=Primates+Dermoptera; Mason et al., 2016) entirely. Definitive stem-euarchontans older than *P. mckeeveri* have not been identified. The oldest members of the sister-taxon of Euarchonta, namely Glires, are younger than the oldest known material of *P. mckeeveri* (see “Crown Glires” below). Thus, *P. mckeeveri* is the oldest known taxon that can be used to calibrate the Euarchonta-Glires split.

<u>Hard minimum bound</u> 65.79 Ma

<u>Soft maximum bound</u> 125.816 Ma

<u>Suggested prior distribution</u> offset exponential

<u>Age justifications</u> High resolution geochronological data constrains the age of the oldest known material of *P. mckeeveri* to the early Puercan (Pu1), chron C29r, and specifically to within ∼208 kyr after the K/Pg boundary (Wilson Mantilla et al., 2021). Wilson Mantilla et al. (2021) reported an 40Ar/39Ar data of a tuff 78 cm located above the Harley’s Point locality, source of our calibrating specimen UCMP 150021, of 65.844 ± 0.033/0.054 Ma (with the uncertainty shown as analytical/systematic uncertainty). We thus use a minimum age of 65.79 Ma for this node. A second date from an underlying tuff has an age of 66.052 ± 0.008/0.043 Ma, bracketing the age of UCMP 150021 to 66.095-65.79 Ma; we use the maximum age as our soft maximum bound for the age of crown Primates (see below).

Placing a maximum bound on the age of Euarchontoglires is difficult because early members of Placentalia, including stem members of Euarchontoglires, probably differed little from stem-eutherians in terms of their overall morphology (Bininda-Emonds et al., 2012). This may explain why probable early placentals such as *Purgatorius* (generally accepted as a euarchontan; see above) and *Protungulatum* (often placed within Laurasiatheria, typically close to euungulates; O’Leary et al., 2013) fall outside Placentalia in some phylogenetic analyses (e.g., Wible et al., 2007, 2009; Goswami et al., 2011). We have chosen to use a conservative maximum bound based on the age of well preserved eutherians from the Yixian Formation such as *Ambolestes*, *Eomaia*, and *Sinodelphys*, which have fallen outside Placentalia in all published phylogenetic analyses to date (e.g., Ji et al., 2002; Kielan-Jaworowska et al., 2004; Bi et al., 2018). The age of the Yixian Formation has now been constrained to between 125.755 ± 0.061 and 124.122 ± 0.048 Ma, based on U-Pb chemical abrasion-isotope dilution-isotope ratio mass spectrometry (CA-ID-IRMS; Zhong et al., 2021); we use the maximum age of this range (125.816 Ma) as our maximum bound here. However, this is almost certainly highly conservative, as the oldest convincing records of Placentalia are from the earliest Palaeocene (e.g., *Purgatorius mckeeveri*; Wilson Mantilla et al., 2021) or very slightly before (e.g., the latest Cretaceous *Protungulatum coombsi*; Archibald et al., 2011). The putative leptictidan *Gypsonictops* is known from the late Cretaceous (Campanian-Maastrichtian; Kielan-Jaworowska et al., 2004) but has also been reported from much older, Turonian deposits (Cohen and Cifelli, 2015), although this material remains to be formally published; if *Gypsonictops* is indeed a leptictidan, and if leptictidans are crown clade placentals, then this would push the origin of Placentalia considerably earlier than the K-Pg boundary, but this remains uncertain (see Springer et al., 2019; Marjanović, 2021). The preponderance of fossil evidence, however, appears to support an origin of Placentalia and of the major placental superorders closer to the K-Pg boundary, and we explicitly take this into account by proposing that this calibration be modelled as an offset exponential prior distribution. Assuming a 5% probability of exceeding the soft maximum bound, this would give a mean and median prior on this divergence of 85.9 and 79.7 Ma, respectively.

<u>Additional CladeAge calibration</u> As discussed above, *Purgatorius mckeeveri* is the oldest known member of Euarchonta. The sister-taxon of Euarchonta is Glires, and the oldest definitive members of Glires (*Heomys* sp., *Mimotona wana*, and *M. lii*) are from the lower part of the Upper Member of the Wanghudun Formation in Anhui Province, China (Li, 1977; see “Crown Glires” below). The lower part of the Upper Member of the Wanghudun Formation is currently interpreted as the Shanguan Stage spanning 66.0-62.22 Ma (Y. Wang et al., 2019). Anagalidans have been suggested to be stem-members of Glires (López-Torres and Fostowicz-Frelik, 2018), and the earliest members of this group are from the Lower Member of the Wanghudun Formation (Marjanović, 2021: node 155), thus predating *Heomys* sp., *Mimotona wana*, and *M. lii*. However, the precise relationship of Anagalida to Glires remains to be clearly resolved (López-Torres and Fostowicz-Frelik, 2018), and we note that they fell outside Euarchontoglires in the recent phylogenetic analysis of Asher et al. (2019). In addition, Wang et al. (2019) did not provide separate ages for the Lower Member and the lower part of the Upper Member of the Wanghudun Formation, and so the same age range (66.0-62.22 Ma) would apply even if we elected to use anagalidans for this additional CladeAge calibration.

<u>Comments</u> The material of *Purgatorius mckeeveri* recently described by Wilson Mantilla et al. (2021) results in a very slightly older minimum on the age of this node than assumed in some previous studies (Benton et al., 2015; Marjanović, 2021). Benton et al. (2009) followed a similar approach to that used here, and based their soft maximum bound on the age of the Yixian eutherians, which were the oldest definitive eutherians known at the time. In their revised list of calibrations, Benton et al. (2015) used the maximum age of an even older eutherian discovered subsequently, namely *Juramaia sinensis* (Luo et al., 2011) to set a maximum bound of 164.6 Ma; however, such a liberal maximum bound is unlikely to place much constraint on the age of this node. By contrast, Marjanović (2021: node 152) proposed a hard maximum bound of 72 Ma on the age of Placentalia, which would force the maximum age of Euarchontoglires (and all other divergences within Placentalia) to be younger than this; however, this very tight maximum bound implicitly endorses an “explosive” model of placental origins, and the validity of this model remains controversial (Springer et al., 2019). We prefer to leave this node quite loosely calibrated, (although not as loosely as Benton et al., 2015), an approach that we feel is warranted given continuing uncertainty regarding the timing of the origin of Placentalia (Bininda-Emonds et al., 2012; Foley et al., 2016; Springer et al., 2019).

### 5.2 Crown Glires = Rodentia-Lagomorpha split

Calibrating taxon *Heomys* sp.

<u>Specimen</u> IVPP V4323, a crushed skull without cheek teeth, from the lower part of the Upper

Member of the Wanghudun Formation in Anhui Province, China (Li, 1977).

<u>Phylogenetic justification</u> Recent phylogenetic analyses (Asher et al., 2019; Rankin et al., 2020) support *Heomys* sp. as the earliest known member of Simplicidentata, which includes Rodentia. Of perhaps greatest significance, simplicidentates (including *Heomys* sp.) are characterized by the presence of a single pair of upper incisors, a synapomorphic feature shared by all rodents (Li et al., 2016; Fostowicz-Frelik, 2017, 2020).

<u>Hard minimum</u> bound 62.22 Ma

<u>Soft maximum</u> bound 66 Ma

<u>Suggested prior distribution</u> offset exponential

<u>Age justifications</u> IVPP V4323 is from the lower part of the Upper Member of the Wanghudun Formation in Qianshan, Anhui Province (Li, 1977). According to Wang et al. (2019), the Lower Member and lower part of the Upper Member of the Wanghudun Formation can be correlated to the Shanghuan Stage, corresponding to chron C27n to C29r, which is 66.0 to 62.22 Ma; the latter date therefore provides the hard minimum bound for this node.

As summarized by Marjanović (2021: node 155), crown-clade Glires (rodents and other simplicidentates; lagomorphs and other duplicidentates) have not been found in the earlier, Lower Member of the Wanghudun Formation (Wang et al., 2016). However, fossil sites from the Lower Member are characterized by a diverse range of anagalidans (anagalids, the pseudictopid *Cartictops* and the astigalid *Astigale*; Marjanović, 2021: node 155). The affinities of anagalidans are in need of detailed study, but it is widely accepted that they include stem relatives of crown Glires (Meng et al., 2003; Fostowicz-Frelik, 2017; López-Torres and Fostowicz-Frelik, 2018; but see the phylogenetic analysis of Asher et al., 2019). Evidence from their molar structure (including a tendency to hypsodonty) and tooth wear suggests that anagalidans were at least partially herbivorous (Fostowicz-Frelik, 2017), as is also the case for most living and fossil members of crown Glires, and features of the postcranial skeleton indicates that pseudictopids were cursorially adapted, similar to lagomorphs (Rose, 2006). We consider the presence in the Lower Member of the Wanghudun Formation of probable stem Glires (namely anagalidans, including some that were probably ecologically similar to lagomorphs), in combination with the apparent absence of crown Glires, to be reasonable evidence that the Rodentia-Lagomorpha split had not occurred by this time. We therefore propose the maximum age of the Shanghuan Stage (66.0 Ma; Y. Wang et al., 2019) as the soft maximum bound on this node.

Given our assumption that the absence of crown Glires in the Lower Member of the Wanghudun Formation is not an artefact of incomplete sampling, but that it is an indication that Rodentia-Lagomorpha split had yet to occur, we consider that this calibration is most appropriately modelled using an offset exponential distribution. Assuming a 5% probability of exceeding the soft maximum bound, this would give a mean and median prior on this divergence of 63.5 and 63.1 Ma respectively.

<u>Additional CladeAge calibration</u> Our calibrating taxon, *Heomys* sp., is the oldest known stem-rodent. The duplicidentates *Mimotona wana* and *M. lii* are from the same deposit as *Heomys* sp.(Li, 1977; Dashzeveg and Russell, 1988; Li et al., 2016). *Mimotona* and other duplicidentates differ from simplicidentates such as rodents and *Heomys* sp. but resemble lagomorphs in retaining two upper incisors (Li et al., 2016; Fostowicz-Frelik, 2020). However, presence of two upper incisors is plesiomorphic for crown Glires, and so does not by itself constitute evidence that duplicidentates are stem-lagomorphs rather than stem-Glires. Nevertheless, *Mimotona* does share one distinctive derived dental synapomorphy with lagomorphs that is not seen in rodents or other simplicidentates, namely a longitudinal groove on the labial surface of the anteriormost upper incisor (Li and Ting, 1993; Li et al., 2016). *Mimotona* was also placed as a stem-lagomorph in the recent phylogenetic analyses of Asher et al. (2019) and Rankin et al. (2020). Based on this collective evidence, we consider *Mimotona wana* and *M. lii* the oldest known stem representatives of Lagomorpha, at 66.0-62.22 Ma, and so provide the second CladeAge calibration for this node.

<u>Comments</u> Benton et al. (2009) proposed a similar minimum bound for this node as that proposed here, but they proposed a much older maximum bound of 131.5 Ma, based on the maximum age of stem eutherians from the Yixian Formation (see “Crown Euarchontoglires” above). Benton et al. (2015) proposed an even older maximum bound, 164.6 Ma, based on the maximum age of the oldest currently known stem eutherian *Juramaia* (see “Crown Euarchontoglires” above). However, we consider both of these maximum bounds to be unduly conservative given the distinctive craniodental apomorphies of members of crown-Glires, and the failure to find such taxa in any mammal-bearing site from the Cretaceous (see also Marjanović, 2021: node 155).

### 5.3 Crown Euarchonta = Scandentia-Primatomorpha split

<u>Calibrating taxon</u> Purgatorius mckeeveri

<u>Specimen</u> UCMP 150021, an isolated lower m2, from Harley’s Point locality in the lowermost part of the Tullock Member of the Fort Union Formation in Montana, USA (Wilson Mantilla et al., 2021).

<u>Phylogenetic justification</u> Retrotransposon insertions provide statistically significant support for the hypothesis that Primates and Dermoptera form a clade (= Primatomorpha) to the exclusion of Scandentia (Mason et al., 2016). Some published analyses examining deep relationships within Eutheria have recovered *Purgatorius* outside Placentalia (e.g., Wible et al., 2007, 2009; Goswami et al., 2011), but all recent published phylogenetic analyses focused on euarchontan relationships have placed *Purgatorius* closer to Primates and/or Dermoptera than Scandentia (Bloch et al., 2007; Ni et al., 2013, 2016; Chester et al., 2015, 2017; Li and Ni, 2016; Gunnell et al., 2018; Morse et al., 2019; Seiffert et al., 2020; see “Crown Euarchontoglires” above).

<u>Hard minimum</u> bound 65.79 Ma

<u>Soft maximum</u> bound 125.816 Ma

<u>Suggested prior distribution</u> offset exponential

<u>Age justifications</u> In contrast to the craniodentally distinctive early crown members of Glires such as *Heomys* and *Mimotona* (see “Crown Glires” above), the earliest crown euarchontans may have been morphologically little different from stem-eutherians (Bininda-Emonds et al., 2012), which might explain why, for example, *Purgatorius* falls outside Placentalia in some published analyses (Wible et al., 2007, 2009; Goswami et al., 2011). For this reason, we use the same minimum and maximum bounds for this node as for Crown Euarchontoglires, and again suggest modelling this as an offset exponential prior (see “Crown Euarchontoglires” above).

<u>Additional CladeAge calibration</u> The fossil record of Scandentia is sparse. Besides the questionable *Eodendrogale parva* from the middle Eocene (Tong, 1988; Ni and Qiu, 2012), the oldest known scandentian is *Ptilocercus kylin* from the early Oligocene Lijiawa locality, Yunnan Province, China, which has an age estimate of ∼34 Ma (Li and Ni, 2016), and represents a second CladeAge calibration for this node. Phylogenetic analyses place *P. kylin* within crown Scandentia, sister to the extant *P. lowii*, suggesting an extensive unsampled history of earlier scandentians.

<u>Comments</u> Benton et al. (2009) proposed similar minimum and maximum bounds to those used here, whilst Benton et al. (2015) instead suggested a maximum bound of 164.6 Ma based on the maximum age of the oldest currently known stem eutherian, *Juramaia* (see “Crown Euarchontoglires” above). Marjanović (2021) did not calibrate this node.

### 5.4 Crown Primatomorpha = Primates-Dermoptera split

<u>Calibrating taxon</u> Teilhardina brandti

<u>Specimen</u> UM 99301 (holotype), an isolated m2, from UM locality SC-351 at the head of Big Sand Coulee in the Clarks Fork Basin, Wyoming (Gingerich, 1993a).

<u>Phylogenetic justification</u> As summarized above (see “Crown Euarchontoglires” above), the precise relationships of the various “plesiadapiforms” to the extant primatomorphan orders Primates and Dermoptera differ quite markedly between recent phylogenetic analyses; we therefore consider them unsuitable for calibrating this node. Two groups with Paleocene representatives, namely plagiomenids and mixodectids, have been proposed by some authors to be dermopteran relatives (Rose, 2006), but this has been questioned (MacPhee et al., 1989; Yapuncich et al., 2011), and so we have also chosen not to use these to calibrate this node. Instead, we use the oldest well-supported member of crown Primates, the omomyiform *Teilhardina brandti* (Gingerich, 1993a; Rose et al., 2011; Morse et al., 2019) as a necessarily conservative minimum bound.

<u>Hard minimum bound</u> 55.935 Ma

<u>Soft maximum bound</u> none

<u>Suggested prior distribution</u> not applicable (minimum bound only)

<u>Age justifications</u> The oldest known material of *Teilhardina brandti*, including our calibrating specimen UM 99301, comes from the Bighorn Basin in Wyoming (Gingerich, 1993a; Smith et al., 2006; Rose et al., 2011; Morse et al., 2019). *T. brandti* material has been reported from various localities in the Bighorn Basin: Big Sand Coulee in the Clarks Fork Basin, a northern sub-basin in the Bighorn Basin (Gingerich, 1993a; Smith et al., 2006), the Willwood formation (Bown and Rose, 1987), and the Sand Creek Divide and Cabin Fork sections (Rose et al., 2011; Morse et al., 2019). All these localities correlate to the second earliest biozone of the early Eocene, Wasatchian-0 (Wa-0), which follows the brief Wa-M biozone and coincides with most of the Paleocene-Eocene Thermal Maximum (PETM; Rose et al., 2011) which is marked by a global carbon isotope excursion (CIE; (Yans et al., 2006). Rose et al. (2011) reported that, based on carbon isotopic stratigraphy, *Teilhardina brandti* appeared only 25 kyr after the onset of the PETM. Using 56.01 ± 0.05 Ma for the start of the PETM (Zeebe and Lourens, 2019), the age estimate of the appearance of *Teilhardina brandti* 25 kyr after this is then 55.985 ± 0.05 Ma, giving a minimum age of 55.935 Ma for UM 99301, which we use as our minimum bound here, and a maximum age of 56.035 Ma.

If the early Palaeocene *Purgatorius* is closer to Primates than to Dermoptera, or vice versa, then it seems likely that the Primates-Dermoptera split could predate the K-Pg boundary; conversely, if *Purgatorius* and other early “plesiadapiforms” are stem rather than crown primatomorphans, then the Primates-Dermoptera split could potentially be close to the Palaeocene-Eocene boundary (the age of the oldest definitive, crown, primates). For this reason, we have chosen not to propose a soft maximum bound on this node.

<u>Additional CladeAge calibration</u> the oldest known definitive dermopteran material is Dermoptera indet. from the Pondaung Formation of Myanmar (Marivaux et al., 2006a), which has been radiometrically dated to 40.31-40.22 Ma ((Khin Zaw et al., 2014; Jaeger et al., 2019).

<u>Comments:</u> This node does not appear to have been calibrated in recent molecular clock analyses, perhaps because compelling evidence for monophyly of Primatomorpha has only become available comparatively recently (Mason et al., 2016).

### 5.5 Crown Primates = Haplorhini-Strepsirrhini split

<u>Calibrating taxon</u> Teilhardina brandti

<u>Specimen</u> UM 99301 (holotype), an isolated m2, from UM locality SC-351 at the head of Big

Sand Coulee in the Clarks Fork Basin, northwestern Wyoming (Gingerich, 1993a).

<u>Phylogenetic justification</u> *Teilhardina brandti* has been identified as an omomyiform (Gingerich, 1993a; Rose et al., 2011; Morse et al., 2019). Phylogenetic analyses consistently place Omomyiformes generally, and *Teilhardina* specifically, within crown Primates. In these analyses, omomyiforms are usually placed within Haplorhini as stem-members of the lineage leading to modern tarsiers (=Tarsiiformes), with which they share large orbit size, elongated tarsals, small body size, an anteriorly positioned foramen magnum indicating a vertical head posture, and shortened crania (Ni et al., 2013, 2016; Gunnell et al., 2018; Jaeger et al., 2019; Morse et al., 2019). Even if *T. brandti* and other omomyiforms are discounted as the oldest crown primates (Godinot, 2015, 2017; Gunnell and Miller, 2018), the oldest known stem strepsirrhine (the 55.8-55.12 Ma old *Donrussellia provincialis*; see below) is only slightly younger than the oldest material of *T. brandti*, and so this would have little impact on the minimum bound of this calibration.

<u>Hard minimum</u> bound 55.935 Ma

<u>Soft maximum</u> bound 66.095 Ma

<u>Suggested prior distribution</u> offset exponential

<u>Age justifications</u> The minimum bound is based on the minimum age of the oldest specimen of the oldest crown primate, *Teilhardina brandti* (see “Crown Primatomorpha” above). The maximum bound is based on the maximum age of the oldest specimen of the oldest known plesiadapiform *Purgatorius mckeeveri* (see “Crown Euarchontoglires” above). Although the affinities of *Purgatorius* and other “plesiadapiforms” vary between analyses, they have not been recovered within crown Primates in any recently published study of which we are aware. A diversity of “plesiadapiforms” is known throughout the Palaeocene (Silcox et al., 2017). At least some of them were likely ecologically similar to early crown primates (Silcox et al., 2017), and they are known from fossil deposits in the same regions (particularly North America) where crown primates are known from younger sites. The presence of “plesiadapiforms” but the absence of ecologically similar crown primates in these Palaeocene sites (several of which are comparatively rich and well-sampled), and the approximately synchronous appearance of members of Haplorhini (*Teilhardina* spp.) and Strepsirrhini (*Donrussellia* spp.) in the earliest Eocene, collectively suggests to us that crown Primates probably originated close to the Palaeocene-Eocene boundary. Based on this, we suggest that this calibration is most appropriately modelled as an offset exponential prior. Assuming a 5% probability of exceeding the soft maximum bound, this would give a mean and median prior on this divergence of 59.3 and 58.3 Ma respectively.

<u>Additional CladeAge calibration</u> We recognize *Teilhardina brandti* as the oldest known haplorhine. Based on current evidence, the oldest known strepsirrhine appears to be the adapiform *Donrussellia*, with three species known from various early Eocene (MP7) sites in Europe (*Donrussellia magna* and *D. provincialis* from France: Godinot, 1978, 1998; and *D. lusitanica* from Portugal: Estravís, 2000). A fourth species, *D. gallica*, is slightly younger (MP8+9; Ramdarshan et al., 2015). Of these, *Donrussellia provincialis* and *D. gallica* have had their phylogenetic affinities formally tested in the context of large scale analyses (e.g., Ni et al., 2013; Morse et al., 2019), and are usually recovered as stem strepsirrhines. *Donrussellia provincialis* is also the best known species, based on multiple dental specimens and an isolated astragalus from the Rians locality (Boyer et al., 2017). We therefore use *D. provincialis* as our additional CladeAge calibration as the oldest known strepsirrhine, with an age range of 55.8-55.12 Ma, based on Solé et al.’s (2015) suggested age for Rians.

<u>Comments</u> We differ from Benton et al. (2015) and dos Reis et al. (2018), who used *Altiatlasius koulchii* from the Adrar Mgorn 1 locality, Morocco (Sigé et al., 1990), as the earliest record of crown primates. Adrar Mgorn 1 can be correlated to Chron 24r (Seiffert et al., 2010b), which spans the Paleocene-Eocene boundary, but based on associated fauna of invertebrates and selachians a latest Paleocene age for Adrar Mgorn 1 appears more likely (Gheerbrant, 1998; Seiffert et al., 2010b). This results in a minimum age of 56.0 Ma for *Altiatlasius koulchii*, based on the age of the end of the Thanetian (Walker et al., 2018), which is only 0.065 Ma older than the minimum bound we set based on the appearance of *Teilhardina brandti*. However*, Altiatlasius* is of very uncertain phylogenetic relationships: it has been identified as a stem primate (Hooker et al., 1999; Morse et al., 2019), a crown primate of uncertain affinities (Silcox, 2008), a stem tarsiiform (Boyer et al., 2010), a basal haplorhine (Marivaux, 2006; Patel et al., 2012), or a stem anthropoid (Godinot, 1994; Marivaux, 2006; Bajpai et al., 2008; Seiffert et al., 2009; Tabuce et al., 2009; Patel et al., 2012) by different authors. Additionally, Seiffert et al. (2010b) note that the morphological variation shown by the upper molars of *Altiatlasius* is problematic, although they still conclude that *Altiatlasius* is more likely to be an anthropoid than a plesiadapiform. Given the uncertainty surrounding its relationships, and the fact its minimum age being very close to that of *Teilhardina brandti*, we do not use *A. koulchii* to calibrate this node.

Benton et al. (2015) used a similar maximum bound to ours, but dos Reis et al. (2018) preferred a much older maximum (88.6 Ma), based on the results of statistical modelling of primate diversification by Wilkinson et al. (2011). In principle, such a quantitative approach is preferable to the admittedly subjective interpretation of the fossil record used here and in most other attempts to identify fossil calibrations for primates. However, we consider a Cretaceous origin for crown Primates to be highly unlikely. Not only is there no record of crown Primates from any Cretaceous site, including the comparatively well-sampled North American record (Kielan-Jaworowska et al., 2004; Wilson, 2014), but a Cretaceous origin for crown Primates would require that all deeper nodes within Euarchontoglires also occurred in the Cretaceous or earlier; there is, however, no record of Cretaceous “plesiadapiforms” either. Furthermore, there does not appear to be a clear explanation why the plesiomorphic “plesiadapiform” *Purgatorius* (a small-bodied [∼100g], predominantly insectivorous, arboreal form; Chester et al., 2015; Silcox et al., 2017; Wilson Mantilla et al., 2021) should appear in the fossil record almost immediately after the K-Pg boundary (Wilson Mantilla et al., 2021), but the oldest crown primates, which appear to have been ecologically broadly similar to *Purgatorius*, appear ∼10 Ma later (and approximately simultaneously in North America, Asia and Europe; Smith et al., 2006; Beard, 2008; Rose et al., 2011) if the lineages leading to *Purgatorius* and crown primates had already diverged in the Cretaceous. Instead, we consider the late Cretaceous and Paleocene fossil record to be sufficiently well sampled to support an origin of crown Primates close to the Palaeocene-Eocene boundary.

### 5.6 Crown Strepsirrhini = Lorisiformes-(Lemuriformes+Chiromyiformes) split

<u>Calibrating taxon</u> Saharagalago misrensis

<u>Specimen</u> CGM 40266 (type), a lower first molar from the BQ-2 locality in the Fayum region,

Egypt (Seiffert et al., 2003).

<u>Phylogenetic justification</u> Recent phylogenetic analyses of *Saharagalago misrensis* consistently place it as a crown strepsirrhine, typically as a stem lorisiform (Gunnell et al., 2018; Seiffert et al., 2018, 2020). A second taxon from BQ-2, *Karanisia clarki*, is also usually placed as a crown strepsirrhine (Gunnell et al., 2018; Seiffert et al., 2018, 2020; López-Torres and Silcox, 2020), providing further evidence that the Lorisiformes-(Lemuriformes+Chiromyiformes) split predates the age of this locality.

<u>Hard minimum bound</u> 37.0 Ma

<u>Soft maximum bound</u> 55.8 Ma

<u>Suggested prior distribution</u> uniform

<u>Age justifications</u> The BQ-2 locality has been correlated with Chron 17n.1n (Seiffert, 2006), which is currently recognized as spanning ∼37.5-37.0 Ma (Houben et al., 2019; Agnini et al., 2020), resulting in a minimum bound for this node of 37.0 Ma (see also Van Couvering and Delson, 2020). The maximum bound is based on the maximum age of the earliest well known stem strepsirrhine, *Donrussellia provincialis* (see “Crown Primates” above), based on the assumption that the divergence of crown Strepsirrhini is unlikely to predate the oldest stem member of the clade. There is a comparatively rich record of stem strepsirrhines from the Eocene in Europe, but the African record is still poorly known, with only three definitive stem strepsirrhines (*Djebelemur*, *Azibius* and *Algeripithecus*) known from the middle Eocene (∼48 Ma; Van Couvering and Delson, 2020) of Algeria and Libya (Tabuce et al., 2009; Marivaux et al., 2013), followed by a ∼11 million year gap until the probable crown strepsirrhines *Saharagalago* and *Karanisia* from BQ-2 mentioned above. For this reason, we suggest that this calibration be implemented as a uniform prior between the minimum and maximum bounds.

<u>Additional CladeAge calibration</u> We accept *Saharagalago* as the oldest known lorisiform. We consider the oldest well supported member of the sister-taxon of Lorisiformes, namely the Chiromyiformes+Lemuriformes clade, to be the stem chiromyiform *Plesiopithecus teras*, from Quarry L-41 in the Fayum region, Egypt (Simons, 1992; Gunnell et al., 2018), which is dated to ∼33.4 Ma (Van Couvering and Delson, 2020; see “Chiromyiformes-Lemuriformes split” below,).

<u>Comments</u> While we have used *Saharagalago* to provide the minimum bound on this node, Benton et al. (2015) instead used *Karanisia clarki* (Seiffert et al., 2003), to provide a minimum bound on the age of this node. In its original description, *Karanisia* was placed as a crown lorisid (Seiffert et al., 2003), but its position in subsequent studies has varied, having been found as a stem-strepsirrhine, stem lorisiform, or stem lemuriform (see summary in López-Torres and Silcox, 2020). We therefore prefer to use *Saharagalago*, which has been consistently placed as a lorisiform in recent analyses, to calibrate this node, as did dos Reis et al. (2018). For the maximum bound, both Benton et al. (2015) and dos Reis (2018) used the age of *Altiatlasius*, which they recognized as the oldest crown primate. However, as already discussed (see “Crown Primates” above), *Altiatlasius* is of uncertain affinities, and we instead use the age of the early stem strepsirrhine, *Donrussellia provincialis*, as our maximum bound here. Nevertheless, the ages of the minimum and maximum bounds proposed here fall closely to those of Benton et al. (2015) and dos Reis et al. (2018).

### 5.7 Chiromyiformes-Lemuriformes split

<u>Calibrating taxon</u> Plesiopithecus teras

<u>Specimen</u> DPC 12393, a crushed but nearly complete cranium with maxillary dentition from Quarry L-41 in the Fayum region, Egypt (Simons, 1992; Simons and Rasmussen, 1994).

<u>Phylogenetic justification</u> Gunnell et al. (2018) presented compelling morphological evidence that *Plesiopithecus* (and a second taxon, the Miocene *Propotto*) is a stem member of Chiromyiformes (see also comments by Godinot, 2006), which today is represented by a single species, the aye-aye *Daubentonia madagascariensis*. This conclusion is supported by total evidence dating phylogenetic analyses, with and without the use of a clock model (Gunnell et al., 2018).

Hard minimum bound 33.4 Ma

Soft maximum bound 55.8 Ma

Suggested prior distribution uniform

<u>Age justifications</u> *Plesiopithecus teras* comes from Quarry L-41 in the Fayum region, Egypt. The age of L-41 has been debated (Gingerich, 1993b; Seiffert, 2006, 2010; see summary in Van Couvering and Delson, 2020), but we follow Van Couvering and Delson (2020) in assigning it an age of 33.4 Ma.

Our proposed maximum bound is the same as for crown Strepsirrhini (see above). In particular, *Plesiopithecus* shows a range of unusual chiromyiform specializations (Godinot, 2006; Gunnell et al., 2018), suggesting that it probably postdates the Chiromyiformes-Lemuriformes split quite considerably, and implying an extensive unsampled ghost lineage. The Oligocene *Bugtilemur mathesoni* from the Bugti Hills, Pakistan, was originally described as a crown lemuriform (Marivaux et al., 2001), but was subsequently identified as an adapiform, and hence a stem strepsirrhine, following the discovery of additional specimens (Marivaux et al., 2006b). The 37.0-37.5 Ma old *Karanisia* was placed as a stem lemuriform in the tip-dating analysis of Seiffert et al. (2018), but most other analyses place it as a stem lorisiform (see summary in López-Torres et al., 2020). Thus, no definitive stem lemuriform fossils are currently known. However, if the Chiromyiformes-Lemuriformes split occurred in mainland Africa, as concluded by Gunnell et al. (2018), then lemuriforms should be expected to be found in the African fossil record. Indeed, Gunnell et al. (2018) implied that the poorly known *Notnamaia* from the middle Eocene (∼47 Ma; Van Couvering and Delson, 2020) of Namibia (Pickford et al., 2008) might be a stem lemuriform (but see Godinot et al., 2018), although this has not (to our knowledge) been tested via formal phylogenetic analysis. Thus it seems possible that the Chiromyiformes-Lemuriformes split might be much older than 33.4 Ma. We therefore consider that a uniform age prior is most appropriate for this node.

<u>Additional CladeAge calibration</u> *Plesiopithecus teras* is the oldest known chiromyiform. The oldest definitive records of the sister clade of Chiromyiformes, Lemuriformes, are subfossil remains from Madagascar, the earliest of which are *Hadropithecus stenognathus,* dating to about 7500 years ago (Burney et al., 2008; Godfrey et al., 2010); this record provides a very young additional CladeAge calibration.

<u>Comments</u> Benton et al. (2015) and dos Reis et al. (2018) did not calibrate this node.

### 5.8 Crown Lorisiformes = Lorisidae-Galagidae split

<u>Calibrating taxon</u> *Komba robustus*

<u>Specimen</u> KNM-SO 501 (holotype), a right mandibular fragment with p4–m2, from Songhor, Kenya (Le Gros Clark and Thomas, 1952).

<u>Phylogenetic justification</u> A position for *Komba* within crown lorisiforms, as a galagid, receives consistently strong support in recent published phylogenetic analyses (Gunnell et al., 2018; Seiffert et al., 2018, 2020). The older *Saharagalago* (see “Crown Strepsirrhini” above) *Karansia* and *Wadilemur* have been recovered as stem galagids in some analyses, but are placed outside crown Lorisiformes in others (see summaries in López-Torres and Silcox, 2020; López-Torres et al., 2020). Also of note is the finding of Phillips (2016) and Phillips and Fruciano (2018) that use of *Saharagalago* to calibrate the lorisid-galagid split results in extremely high apparent dating error. Using the results of molecular dating analyses to assess the appropriateness of particular fossil calibrations risks circularity, but in this case the strong mismatch in molecular rates found by Phillips (2016) and Phillips and Fruciano (2018) when *Saharagalago* is assumed to be a crown lorisiform, together with the fact that *Saharagalago* falls outside crown Lorisiformes in at least some analyses (Gunnell et al., 2018; Seiffert et al., 2018, 2020), persuades us that *Komba robustus* is a more appropriate calibrating fossil taxon for this divergence.

<u>Hard minimum bound</u> 18.5 Ma

<u>Soft maximum bound</u> 55.8 Ma

<u>Suggested prior distribution</u> uniform

<u>Age justifications</u> species of *Komba* (as well as several other putative galagids that have not had their phylogenetic affinities robustly tested, such as *Progalago* spp. and *Mioeuoticus* spp.) are known from multiple early Miocene sites in east Africa (Harrison, 2010a: table 20.2). Although radiometric dates are available for at least some of these sites, including Songhor from where KNM-SO 501 was collected, this dating was done in the 1960s using K-Ar dating (Bishop et al., 1969), and it is in need of verification using more modern techniques (Cote et al., 2018). Songhor is currently recognized as falling within the Legetetian African Land Mammal Age (Van Couvering and Delson, 2020), and so pending new radiometric dating of this site, we use the minimum age of the Legetetian (which spans 22.5-18.5 Ma according to Van Couvering and Delson, 2020) as our minimum bound here, namely 18.5 Ma. Our maximum bound is the same as for crown Strepsirrhini and Chiromyiformes-Lemuriformes (see above).

Given that *Saharagalago*, *Karanisia* and *Wadilemur* have all been recovered as stem galagids in some analyses, it is possible that the galagid-lorisid split predates considerably our proposed minimum bound. For this reason, this calibration is most appropriately modelled as a uniform distribution.

<u>Additional CladeAge calibration</u> The oldest lorisid that has had its phylogenetic affinities rigorously tested is *Nycticeboides simpsoni*, which is dated to ∼8.9 Ma, and falls within crown Lorisidae in most recent analyses (see “Crown Lorisidae” below).

<u>Comments</u> Benton et al. (2015) did not calibrate this node. By contrast, dos Reis et al. (2018) used a similar minimum bound to ours (18 Ma), based on the early Miocene *Mioeuoticus*, which they recognized as a crown lorisid, but a tighter maximum bound (38 Ma) that seems questionable given the possibility that the 37.5-37.0 Ma *Saharagalago* is a crown lorisiform (see above); indeed, we note that their 95% posterior credibility interval for the Lorisidae-Galagidae split (34.1-40.9 Ma) exceeds their proposed maximum bound.

### 5.9 Crown Lorisidae = Lorisinae-Perodicticinae split

<u>Calibrating taxon</u> Nycticeboides simpsoni

<u>Specimen</u> YGSP 8091 (holotype), a near complete dentition formed by mandibular and maxillary fragments, some skull fragments, and a few postcranial fragments including a distal humerus, all believed to represent a single individual, from the YGSP 363 locality in the Dhok Pathan Formation, Pakistan (Jacobs, 1981).

<u>Phylogenetic justification</u> *Nycticeboides simpsoni* closely resembles extant *Nycticebus* species (Jacobs, 1981; MacPhee and Jacobs, 1986; Flynn and Morgan, 2005) and is typically found to be a crown lorisine in published phylogenetic analyses: either sister to *Nycticebus* (Seiffert et al., 2015, 2018; Herrera and Dávalos, 2016), or sister to *Loris* (Seiffert et al., 2018). In a few analyses, however, *Nycticeboides* is place as a stem lorisine, outside *Loris*+*Nycticebus* (Seiffert et al., 2015), or as part of an unresolved polytomy with *Loris* and *Nycticebus* (Seiffert et al., 2010a). Regardless, all of these phylogenetic placements support the use of *Nycticeboides* to place the minimum bound for the divergence between Lorisinae and Perodicticinae. An exception to this general pattern is seen in the total-evidence phylogenetic analyses by Seiffert et al. (2018), in which *Nycticeboides* was placed as a stem rather than crown lorisid. However, Gunnell et al. (2018) and Seiffert et al. (2020) both used morphological matrices that were expanded from Seiffert et al. (2018), and in both cases *Nycticeboides* was placed within crown Lorisidae. Morphological synapomorphies that support *Nycticeboides* as a lorisine (and hence a crown lorisid) are found in its facial, dental, and postcranial morphology (Jacobs, 1981; MacPhee and Jacobs, 1986), and so we are confident in using this taxon to calibrate this node here.

<u>Hard minimum bound</u> 8.9 Ma

<u>Soft maximum bound</u> 37.5 Ma

<u>Suggested prior distribution</u> uniform

<u>Age justifications</u> The YGSP 363 locality in the Dhok Pathan Formation, Pakistan, has been argued to be younger than 8 Ma based on dating of older sites in the same section (Tauxe, 1979), and *Nycticeboides* was assigned an approximate age of ∼8-7 Ma in its original description (Jacobs, 1981). MacPhee and Jacobs (1986) listed an age of 7.5-7.0 Ma for the holotype based on tracing of the lithologic unit to a measured section dated by Tauxe and Opdyke (1982). However, Flynn and Morgan (2005) subsequently reported an age of 9.1-7.8 Ma for YGSP 363, and this locality is currently believed to be ∼8.9 Ma old (L. J. Flynn, pers. comm. 21/01/2021); we use this latter date as our hard minimum bound here

As discussed above (see “Crown Strepsirrhini” and ”Crown Lorisiformes” above), most recent published phylogenetic analyses find that the 37.0-37.5 Ma old *Saharagalago* and *Karanisia* are stem lorisiforms, and so it seems likely that they predate divergences within the crown lorisiform families Lorisidae and Galagidae. We therefore use the maximum age of the BQ-2 Quarry (see “Crown Strepsirrhini” above) as our maximum bound here.

There are a number of fossil putative lorisids that are older than *Nycticeboides simpsoni*, at least some of which may be members of crown Lorisidae. These include *Mioeuoticus* from the early Miocene (∼19-18 Ma) of East Africa (Le Gros Le Gros Clark, 1956; Leakey, 1962), ?*Nycticebus linglom* from the Miocene (18.0-17.0 Ma or 14.2-12.0 Ma) of Thailand (Mein and Ginsburg, 1997), and an isolated m1 from the middle Miocene (∼15.2 Ma) locality Y682 in the Kamlial Formation of Pakistan that Flynn and Morgan (2005) identified as *Nycticeboides* sp. We have not used these taxa to inform our proposed minimum bound on this divergence here, because their phylogenetic affinities are either controversial or have not been formally tested; nevertheless, they suggest that the Lorisinae-Perodicticinae split may predate considerably the age of *Nycticeboides simpsoni*, and so a uniform prior distribution on the age of this node seems appropriate.

<u>Additional CladeAge calibration</u> As summarized above, we consider *Nycticeboides simpsoni* to be the oldest well supported member of crown Lorisidae. The affinities of most other fossil lorisids currently known are controversial or have not been tested via formal phylogenetic analysis. Pickford (2012) described OCO 119’10, a partial rostrum (preserving part of the upper dentition) of a lorisid from the Aragai locality in the Lukeino Formation, and tentatively referred this specimen to the extant perodicticine genus *Arctocebus*. Although OCO 119’10 has not been included in a published phylogenetic analysis, its close overall resemblance to *Arctocebus* means that we consider it the oldest definitive perodicticine. The Aragai locality is currently considered to be ∼6.1 Ma (Gilbert et al., 2010).

<u>Comments</u> dos Reis et al. (2018) used a considerably older minimum bound for this divergence of 14 Ma, based on an undescribed genus and species from Fort Ternan in Kenya, which Harrison (2010a) reported “is most similar to *Perodicticus*, and may eventually be referable to the Perodicticinae.” However, pending description of this specimen and formal testing of its affinities, we prefer a younger minimum bound here. The maximum bound of dos Reis et al. (2018) is similar to that used here.

### 5.10 Galagoides*-(*Galago*+*Paragalago*+*Otolemur*+*Sciurocheirus*) split*

<u>Calibrating taxon</u> Galago farafraensis

<u>Specimen</u> SA 8′05, a right p4, from the Sheikh Abdallah locality, Egypt (Pickford et al., 2006b).

<u>Phylogenetic justification</u> *Galago farafraensis* is represented by dental and postcranial material. Although the affinities of this taxon have not been tested via formal phylogenetic analyses, its p4 exhibits a reduced metaconid that is positioned close to the protoconid, a derived trait that is shared with extant *Galago* and *Otolemur* species, in contrast to the plesiomorphic condition in early and middle Miocene galagids and extant *Euoticus* and *Galagoides*, in which the metaconid is larger and more offset from the protoconid (Schwartz and Tattersall, 1985; Walker, 1987; McCrossin, 1992; Pickford et al., 2006b). Although we recognize that this is only a single character, in the absence of a suitably comprehensive phylogenetic analysis, we suggest that *Galago farafraensis* is more closely related to living *Galago, Paragalago, Otolemur,* and *Sciurocheirus* species than to *Euoticus* and *Galagoides* based on this dental synapomorphy, and so *G. farafraensis* can be used to calibrate this divergence. Other fragmentary galagid material from the Miocene and Pliocene has not been formally described, or is so fragmentary that it cannot be confidently placed in a phylogenetic framework, or both (Harrison, 2010a).

<u>Hard minimum bound</u> 10.0 Ma

<u>Soft maximum bound</u> 33.4 Ma

<u>Suggested prior distribution</u> uniform

<u>Age justifications</u> Van Couvering and Delson (2020) assigned an age of ∼11 Ma to Sheikh Abdallah, but we take the more conservative approach of assuming an age range corresponding to the entire Tugenian African Land Mammal Age (12.8-10 Ma), which includes Sheikh Abdallah (Van Couvering and Delson, 2020).

Based on molecular evidence (Pozzi et al., 2014, 2015), this node is the second split within crown Galagidae, with *Euoticus* the first modern galagid genus to diverge. It seems unlikely that a nested divergence within crown Galagidae will predate the oldest fossil record of stem galagids. For this reason, we use the *Wadilemur elegans* from Quarry L-41 in the Fayum, which has been recovered as a stem galagid in most (but not all) recent phylogenetic analyses, to provide our maximum bound (see “Chiromyiformes-Lemuriformes split” above for discussion of the age of Quarry L-41). However, given the fragmentary nature of the galagid fossil record from the Miocene and Pliocene, and the lack of a robust phylogenetic framework for most of these specimens (Harrison, 2010a, 2011; Kunimatsu et al., 2017), it seems possible, if not probable, that this divergence could predate our minimum bound considerably, and so a uniform distribution seems appropriate.

<u>Additional CladeAge calibration</u> An isolated right m2 (L. 1-521) from the upper Member B of the Shungura Formation at Omo, Ethiopia, described by Wesselman (1984) was identified by Harrison (2010a) as *Galagoides* cf. *zanzibaricus* based on its very close resemblance to the modern species. Although this is not the result of a formal phylogenetic analysis, we accept this identification here and use this specimen to provide our additional CladeAge calibration for this node. Van Couvering and Delson (2020) report an age of 3.42 Ma for Member B of the Shungura Formation, and we use this date here.

<u>Comments</u> dos Reis et al. (2018) used a ∼15 Ma old undescribed galagid from Maboko Island, Kenya (McCrossin, 1999), to provide the minimum bound for the calibration of crown Galagidae. We do not calibrate crown Galagidae here, preferring instead to calibrate a less inclusive node. In any case, in the absence of a formal description, it is unclear whether the Maboko Island taxon represents a crown or a stem galagid. For their maximum bound, meanwhile, dos Reis et al. (2018) used a similar, but slightly older date for crown Galagidae to that proposed here for *Galagoides-*(*Galago*+*Paragalago*+*Otolemur*+*Sciurocheirus*), namely 37 Ma based on the age of *Karanisia clarki* from the BQ-2 locality in the Fayum (see “Crown Strepsirrhini” above).

### 5.11 Crown Haplorhini = Anthropoidea-Tarsiiformes split

<u>Calibrating taxon</u> *Tarsius eocaenus*

<u>Specimen</u> IVPP V14563, a left premaxillary-maxillary fragment preserving the crown of P3, alveoli for I2, C1, P2, and the mesial roots of P4, from Shanghuang fissure D, near the village of Shanghuang, southern Jiangsu Province, China (Rossie et al., 2006).

<u>Phylogenetic justification</u> *Tarsius eocaenus* has not, to our knowledge, been included in a comprehensive phylogenetic analysis to formally test its affinities, but its preserved cranial morphology is almost identical to that seen in modern tarsiids, and includes several unusual derived traits (Rossie et al., 2006). Based on this, we are confident that *Tarsius eocaenus* is a definitive tarsiiform. Omomyiforms, including the oldest known member of this group *Teilhardina brandti*, are typically placed as stem tarsiiforms in recent phylogenetic analyses (see “Crown Primates” above). However, some doubts remain as to whether omomyiforms are indeed members of the tarsiiform lineage (Godinot, 2015; Gunnell and Miller, 2018). Based on current evidence, the oldest anthropoids are eosimiids and amphipithecids from the Eocene of Asia (Jaeger et al., 2019, 2020), of which the oldest well dated taxa are from the Pondaung Formation of Myanmar (Zin-Maung-Maung-Thein et al. 2017; Jaeger et al., 2020), which has been radiometrically dated to 40.31-40.22 Ma (Khin Zaw et al., 2014; Jaeger et al., 2019). Older putative records of anthropoids are based on specimens that are much more fragmentary, and are correspondingly more equivocal; they include *Altiatlasius koulchii* from the Palaeocene-Eocene of Africa, which is of very uncertain relationships (see “Crown Primates” above), and *Anthrasimias gujaratensis* from the early Eocene of India (Bajpai et al., 2008), the material of which has subsequently been suggested to in fact represent the asiadapid (stem strepsirrhine) *Marcgodinotius indicus* (Rose et al., 2009, 2018). There is thus a ∼15 million year gap between the oldest omomyiform (*Teilhardina brandti*) and the oldest generally accepted record of stem anthropoids, from the Pondaung Formation. By contrast, *Tarsius eocaenus,* which is from the 47.8-41.0 Ma old (Ni et al., 2020) Shanghuang fissure fills of China, is similar in age to the Pondaung stem anthropoids. While the primate fossil record is obviously far from complete, the large gap between the oldest omomyiforms and the oldest anthropoids may be an indication that omomyiforms are stem rather than crown haplorhines; thus, we prefer to use the *Tarsius eocaenus* to provide a minimum on this node.

<u>Hard minimum bound</u> 41.0 Ma

<u>Soft maximum bound</u> none

<u>Suggested prior distribution</u> not applicable (minimum bound only)

<u>Age justifications</u> Ni et al. (2020) identified the Shanghuang fissure fills as spanning 47.8-41.0 Ma, and so we use 41.0 Ma as our minimum bound. Given the uncertainty regarding the affinities of omomyiforms discussed above, we find it difficult to define a maximum bound, and so we leave this uncalibrated here.

<u>Additional CladeAge calibration</u> We assume that *Tarsius eocaenus* is the oldest definitive member of Tarsiiformes. Similarly, we assume that the oldest definitive stem anthropoids are from the 40.22-40.31 Ma Pondaung Formation (see above), which we use as our additional CladeAge calibration.

<u>Comments</u> Although dos Reis et al. (2018) did not discuss omomyiform affinities, it is notable that they chose to specify the minimum bound on crown Haplorhini using *Tarsius eocaenus* (as done here), together with a second fossil tarsiid from China (*Xanthorhysis*), rather than using an omomyiform. They also, like us, leave the maximum bound on this node uncalibrated.

### 5.12 Crown Anthropoidea = Catarrhini-Platyrrhini split

<u>Calibrating taxon</u> Catopithecus browni

<u>Specimen</u> DPC 8701, a near complete skull, from Quarry L-41 in the Fayum Region, Egypt (Simons, 1989, 1990).

<u>Phylogenetic justification</u> *Catopithecus* can be identified as a stem-catarrhine, and therefore a crown anthropoid, based on the loss of the upper and lower second premolars, and the development of a honing blade for the upper canine on a sexually dimorphic lower p3 (Simons and Rasmussen, 1996; Seiffert and Simons, 2001). The stem catarrhine position of *Catopithecus* is confirmed by numerous recent phylogenetic analyses, including those of Ni et al. (2016), Morse et al. (2019), and Seiffert et al. (2020).

<u>Hard minimum bound</u> 33.4 Ma

<u>Soft maximum bound</u> 56.035 Ma

<u>Suggested prior distribution</u> uniform

<u>Age justifications</u> *Catopithecus browni* comes from Quarry L-41 in the Fayum Region, Egypt, for which we use an age estimate of 33.4 Ma, following Van Couvering and Delson (2020; see “Crown Strepsirrhini” above). For the maximum bound we use the maximum age of our calibrating specimen of the oldest crown primate, *Teilhardina brandti* (see “Crown Primatomorpha” above).

A few stem anthropoids have been described from African sites that are slightly older than Quarry L-41 (e.g., *Biretia*, *Talahpithecus*), but as yet no definitive crown anthropoids; however, a currently undescribed taxon from the 37.5-37.0 Ma BQ-2 locality may represent a stem catarrhine (Gunnell and Miller, 2018; E. R. Seiffert, pers. comm. 24/03/2021), which would result in a slightly older minimum bound than that proposed here. In addition, some phylogenetic analyses presented by Jaeger et al. (2019) placed *Aseanpithecus* from the 40.31-40.22 Ma Pondaung Formation of Myanmar within crown Anthropoidea. For these reasons, at present we consider that this calibration is best modelled as a uniform distribution, although we suspect that this divergence is almost certainly closer to the minimum than the maximum bound.

<u>Additional CladeAge calibration</u> *Catopithecus browni* is the oldest known stem catarrhine. Antoine et al. (2021) recently described highly fragmentary primate teeth from Shapaja, San Martín, Peruvian Amazonia in a site (TAR-21) that they dated to between 33.9 and 34.5 Ma, i.e., the latest Eocene (Antoine et al., 2021, fig. 6). These specimens resemble *Perupithecus ucayaliensis* from the early Oligocene (29.6 ± 0.08 Ma) Santa Rosa Fauna of Peru (Campbell et al., 2021), which is probably a stem platyrrhine (see “Results” and “Discussion” above; Bond et al., 2015; Seiffert et al., 2020), and so we tentatively recognize them as stem platyrrhines as well. However, the reported ages of the Shapaja sites were questioned by Campbell et al. (2021), with these authors concluding that an Oligocene date was more likely. Pending resolution of this issue, we prefer to use the detrital zircon date for the Santa Rosa Fauna, source of the probable platyrrhine *Perupithecus*, as our second CladeAge calibration: this is 29.68-29.52 Ma. *Talahpithecus* from the ∼36 Ma (Van Couvering and Delson, 2020) Dur At-Talah escarpment, central Libya, was recovered as a stem-platyrrhine in the phylogenetic analysis of Bond et al. (2015), but its position as sister to *Perupithecus* implies a very complex biogeographical origin for Platyrrhini with multiple crossings of the Atlantic Ocean, and so we do not use *Talahpithecus* as the oldest record of Platyrrhini here.

<u>Comments</u> Benton et al. (2015) and dos Reis et al. (2018) also used *Catopithecus* to provide a minimum bound on this node. However, both these studies used a more conservative maximum bound than that proposed here. Benton et al. (2015) used 66 Ma, based in part on their identification of *Altiatlasius* as the oldest crown primate and possible crown anthropoid; however, we consider the affinities of *Altiatlasius* to be uncertain (see “Crown Primates” above) and do not use it for calibration purposes. Dos Reis et al. (2018), meanwhile, used a maximum of 62.1 Ma based on the modelling of primate diversification by Wilkinson et al. (2011), about which we have concerns (see “Crown Primates” above).

### 5.13 Crown Catarrhini = Cercopithecoidea-Hominoidea split

<u>Calibrating taxon</u> Rukwapithecus fleaglei

<u>Specimen</u> RRBP 12444A (holotype), a right mandible including p4-m3 and part of ascending ramus from Nsungwe 2B, Tanzania

<u>Phylogenetic justification</u> *Rukwapithecus fleaglei* was consistently recovered as a stem-hominoid (within the clade Nyanzapithecinae) in the parsimony and Bayesian phylogenetic analyses of Stevens et al. (2013), indicating that it postdates the Cercopithecoidea-Hominoidea split. Stevens et al. (2013, fig. 3) noted that some nodes within their illustrated phylogeny have low support values, but there are various synapomorphies reported for four nodes leading up to Nyanzapithecinae, and for this subfamily itself. *Rukwapithecus fleaglei* shares two synapomorphies with Miocene and extant hominoids that are not present in cercopithecoids or stem-catarrhines: a buccal position of the M2 hypoconulid, and the mesial migration of cusps on the buccal side of lower molars such that the hypoconid is positioned opposite the lingual notch between the metaconid and the entoconid (Stevens et al., 2013).

<u>Hard minimum bound</u> 25.193 Ma

<u>Soft maximum bound</u> 33.4 Ma

<u>Suggested prior distribution</u> uniform

<u>Age justifications</u> *Rukwapithecus fleaglei* comes from locality Nsungwe 2B in the Oligocene Nsungwe Formation in southwestern Tanzania (Stevens et al., 2013). The age of the fossil bearing unit is constrained by two volcanic tuffs dated by U-Pb zircon CA-TIMS (U-Pb chemical abrasion thermal ionization mass spectrometry) at 25.237 ± 0.098 and 25.214 ± 0.021 Ma (Stevens et al., 2013). Taking into account these confidence intervals, the minimum age for this specimen is 25.193 Ma and the maximum is 25.335 Ma. For the soft maximum bound, we use the age of the oldest known probable stem-catarrhine, *Catopithecus browni*, from the Quarry L-41 of the Fayum, Egypt (see “Crown Strepsirrhini” above). The late Oligocene record of primates and other terrestrial mammals in Africa is notoriously poor (Kappelman et al., 2003; Wilkinson et al., 2011; Stevens et al., 2013), and for this reason we suggest that this calibration is best modelled as a uniform calibration.

<u>Additional CladeAge calibration</u> Another fossil species from Nsungwe 2B is *Nsungwepithecus gunnelli*, currently known from a single specimen (RRBP 11178), a left partial mandible with a lower m3 (Stevens et al., 2013). *Nsungwepithecus* was not included in the phylogenetic analyses by Stevens et al. (2013), but they report the presence of numerous lower molar synapomorphies that are shared with “victoriapithecid” cercopithecoids (“Victoriapithecidae” is a paraphyletic assemblage of stem cercopithecoids in the phylogenetic analyses of Stevens et al. (2013) and Rasmussen et al. (2019)), such as deeply incised buccal clefts, a high degree of buccal flare, and the lack of a buccal cingulid. Rasmussen et al. (2019) confirmed the stem cercopithecoid position of *Nsungwepithecus* in their phylogenetic analysis, but they argued that the phylogenetic position of *Nsungwepithecus* should be regarded as tentative until more material is available. We therefore recognize *Nsungwepithecus gunnelli* as the oldest (stem) representative of Cercopithecoidea, with the same age as *Rukwapithecus fleaglei*.

<u>Comments</u> Although differing in detail, Benton et al. (2015), dos Reis et al. (2018) and Roos et al. (2019) all proposed very similar minimum and maximum bounds for this node.

### 5.14 Crown Cercopithecidae = Cercopithecinae-Colobinae split

<u>Calibrating taxon</u> Colobinae gen. et. sp. indet.

<u>Specimen</u> KNM-TH 48368, an isolated right lower molar (?m3) from the Baringo Paleontological Research Project (BPRP) no. 38 site in the Kabasero type section of the Ngorora Formation, Tugen Hills succession, Kenya (Rossie et al., 2013).

<u>Phylogenetic justification</u> Phylogenetic analyses by Rossie et al. (2013) consistently placed KNM-TH 48368 as an early colobine, regardless of whether it was coded as an m2 or an m3. KNM-TH 48368 displays a very small but distinct hypoconulid which is also present in the fossil colobines *Microcolobus* and *Mesopithecus* and many extant colobines (Rossie et al., 2013). Synapomorphies that KNM-TH 48368 shares with extant colobines are: “tall and sharp transverse lophids, reduced basal flare of the crown, a wide and deep median buccal cleft, buccal cusps with a columnar profile and mesial tilt, a long talonid basin relative to overall crown length, and subequal mesial and distal crown breadths” (Rossie et al., 2013).

<u>Hard minimum bound</u> 12.47 Ma

<u>Soft maximum bound</u> 25.235 Ma

<u>Suggested prior distribution</u> offset exponential

<u>Age justifications</u> KNM-TH 48368 comes from the Kabasero section of the Ngorora Formation in the Tugen Hills, Kenya (Rossie et al., 2013). 40Ar/39Ar dating of the fossiliferous horizon itself provides an age of 12.49 ± 0.02 Ma for this locality, resulting in a minimum and maximum age for this specimen of 12.47 and 12.51 Ma respectively. The horizon is also bracketed below and above by 40Ar/39Ar dates of 12.56 ± 0.04 Ma and 12.26 ± 0.07 Ma respectively (Deino et al., 2002; Hill et al., 2002; Rossie et al., 2013). The maximum bound for this node is based on the maximum age of the two oldest known crown-catarrhines, namely the stem cercopithecoid *Nsungwepithecus* and stem hominoid *Rukwapithecus* from Nsungwe 2B, Tanzania, with a maximum age of 25.214 ± 0.021 Ma (see “Crown-Catarrhini” above).

Between the oldest known stem cercopithecoid *Nsungwepithecus* and KNM-TH 48368, a diverse range of fossil cercopithecoids are known from multiple early and middle Miocene (∼22.5-15 Ma; Van Couvering and Delson, 2020) sites throughout Africa, comprising at least nine species-level taxa (Locke et al., 2020 table 1). Not all of these have had their phylogenetic affinities formally tested, but those that have (namely the “victoriapithecids” *Prohylobates*, *Noropithecus,* and *Victoriapithecus*) consistently fall outside crown Cercopithecidae (Miller et al., 2009; Stevens et al., 2013; Rasmussen et al., 2019). The African primate fossil record is sparse between 15 and 6 Ma (Rossie et al., 2013). However, the diversity of stem cercopithecids between 22.5 and 15 Ma (Locke et al., 2020 table 1) and the apparent absence of crown cercopithecids in this same time interval persuades us that this divergence is likely to be close to our minimum bound, and so we propose an offset exponential prior distribution. Assuming a 5% probability of exceeding the soft maximum bound, this would give a mean and median prior on this divergence of 16.7 and 15.4 Ma respectively.

<u>Additional CladeAge calibration</u> KNM-TH 48368 is the oldest known colobine. The oldest known record of Cercopithecinae is possible stem papionin material from the Beticha locality in the Chorora Formation, Ethiopia (Suwa et al., 2015; Katoh et al., 2016). Based on available evidence, we do not consider the Beticha material to be unequivocally papionin (see “Crown Cercopithecinae” below), but we do recognize it as the oldest cercopithecine. The Beticha fossil-bearing unit is above a pumiceous tuff that has been dated to 8.18 +/-0.15 Ma by K–Ar dating and 7.86 +/- 0.10 Ma by 40Ar–39Ar dating, and below a consolidated tuff dated to 7.67 +/- 0.17 Ma by K–Ar dating and 7.82 +/- 0.11 Ma by 40Ar–39Ar dating (Katoh et al., 2016). Taking the maximum and minimum bounds for these radiometric dates, this gives an age range of 8.33-7.5 Ma, which we suggest as our additional CladeAge calibration.

<u>Comments</u> dos Reis et al. (2018) do not calibrate this node, but our maximum and minimum bounds are similar to those proposed by Roos et al. (2019).

### 5.15 Crown Colobinae = Colobini-Presbytini split

<u>Calibrating taxon</u> *Mesopithecus pentelicus delsoni* (*Mesopithecus delsoni* according to de Bonis et al., 1990; recognized here as subspecies of *Mesopithecus pentelicus* following Alba et al., 2015).

<u>Specimen</u> RZO 159 (holotype of *Mesopithecus delsoni,* recognized as a subspecies of *Mesopithecus pentelicus* by Alba et al. (2014a, 2015)), a nearly complete adult male mandible, from Ravin des Zouaves-5, Greece.

<u>Phylogenetic justification</u> *Mesopithecus* has been consistently placed as a member of Presbytini in the few published morphological phylogenetic analyses that have specifically examined this question (Jablonski, 1998; Byron, 2001). In addition, dental metrics of *Mesopithecus* are more similar to modern presbytins than to colobins (Pan et al., 2004), and mandibular morphology of *Mesopithecus* shows particular similarities to that of the modern presbytin genera *Rhinopithecus* and *Pygathrix* (Jablonski et al., 2020), although these resemblances are only suggestive because they have not been placed in an explicit phylogenetic context. Some researchers have cited the unreduced pollex of *Mesopithecus* as evidence that it falls outside crown Colobinae, all living members of which are characterized by a reduced-to-absent pollex (with a greater degree of pollicial reduction in colobins than presbytins; Alba et al., 2015; Frost et al., 2015). However, *Mesopithecus* has been reported to have a slightly reduced pollex (but see Frost et al., 2015; Jablonski et al., 2020), and Jablonski (1998: character 148) specifically included “thumb length” as one of the 455 morphological characters used in her phylogenetic analysis. As noted by Jablonski et al. (see also Nakatsukasa et al., 2010; 2020), pollicial reduction has occurred at least twice within Anthropoidea, as the pollex is greatly reduced or absent in the platyrrhine atelids *Ateles* and *Brachyteles* (Rosenberger et al., 2008), and we agree with those authors that undue weight should not be placed on a single morphological character, at least when phylogenetic analyses based on multiple characters are available (Jablonski, 1998; Byron, 2001). We therefore recognize *Mesopithecus* as the earliest definitive presbytin based on the results of available phylogenetic analyses (Jablonski, 1998; Byron, 2001), and therefore suitable for calibrating this node.

<u>Hard minimum bound</u> 8.2 Ma

<u>Soft maximum bound</u> 15 Ma

<u>Suggested prior distribution</u> uniform

<u>Age justifications</u> The source of our calibrating specimen, the Ravin des Zouaves-5 locality in Greece, is estimated to date to ∼8.2 Ma based on magnetostratigraphic evidence, specifically correlation to C4r.1r (Sen et al., 2000; Koufos, 2009), which we therefore use as our minimum bound. A *Mesopithecus* specimen from another Greek locality, Nikiti 2, may slightly pre-date this (Koufos, 2016), but its minimum age is also 8.2 Ma. The material from Nikiti 2 is also very incomplete, comprising one metacarpal and one metatarsal, and we prefer to use the Ravin des Zouaves-5 specimen (which is a near complete mandible) to calibrate this node. A maxillary fragment of *Mesopithecus* has also been reported from Grebeniki 1 (Gremyatskii, 1961), Ukraine, which was originally dated to the early Turolian and the MN11 [8.8-7.9 Ma, following collated information by Alba et al. (2015)], but a recent faunal correlation by Vangengeim and Tesakov (2013) correlated Grebeniki 1 with the preceding MN10 (9.7-8.8 Ma) which would imply an older minimum bound on this node; however, given the current uncertainty surrounding the age of this site (see Koufos, 2019), we do not use it to calibrate this node.

The oldest stem-colobine material is the ∼12.5 Ma Colobinae gen. et. sp. indet. from Tugen Hill (Rossie et al., 2013; see “Crown Cercopithecidae” above). However, we have decided against using this material as the basis for our maximum bound due to the poor African record of primates between 15 and 6 Ma (Rossie et al., 2013); instead, we use 15 Ma as our maximum bound, as the better sampled Miocene record prior to this date reveals a diversity of stem-cercopithecoids (“victoriapithecids”) but no crown forms (Locke et al., 2020: table 1; see “Crown Cercopithecidae” above). Based on this poor record 15-6 Ma, we suggest modelling this calibration as a uniform prior.

<u>Additional CladeAge calibration</u> *Mesopithecus pentelicus delsoni* is the oldest known presbytin. The oldest colobin, meanwhile, appears to be *Cercopithecoides bruneti* from Toros-Menalla, Chad, which can be referred to Colobini rather than Presbytini based on probable synapomorphic traits such as its gracile mandibular morphology and terrestrial locomotion adaptations in its forelimb (Pallas et al., 2019). The dentognathic morphology supports the placement of *Cercopithecoides* as a stem rather than crown colobin (Pallas et al., 2019). The Toros-Menalla locality has been dated to 7 ± 0.2 Ma (Lebatard et al., 2010), giving a CladeAge calibration of 6.8-7.2 Ma.

<u>Comments</u> dos Reis et al. (2018) used the 9.8 Ma colobine *Microcolobus* to provide a minimum bound on this node, but to our knowledge *Microcolobus* has not been demonstrated to be a member of crown Colobinae, and in fact Rossie et al. (2013) found it to be more distantly related to extant colobines than the older Tugen Hills material, suggesting that this taxon is more likely to be a stem form. Their maximum bound of 23 Ma is more conservative than ours, and does not appear to take into account the diverse early Miocene record of stem cercopithecoids (Locke et al., 2020). Roos et al. (2019), meanwhile, used *Mesopithecus* to provide a minimum bound on this node in their “calibration set 1”, as done here, but set their maximum as 12.5 Ma based on the Kabasero colobine material (Rossie et al., 2013; see “Crown Cercopithecidae” above), which, as discussed, we consider overly restrictive given the poor African primate record 15-6 Ma (Rossie et al., 2013).

### 5.16 Crown Cercopithecinae = Cercopithecini-Papionini split

<u>Calibrating taxon</u> Cercopithecini sp. indet

<u>Specimen</u> AUH 1321, a lower left molar, most likely an m1, from the SHU 2-2 locality in the Baynunah Formation, Abu Dhabi (Gilbert et al., 2014).

<u>Phylogenetic justification</u> Published phylogenetic analyses indicate that AUH 1321 is a crown cercopithecin (Gilbert et al., 2014; Plavcan et al., 2019; see “Crown Cercopithecini” below).

<u>Hard minimum bound</u> 6.5 Ma

<u>Soft maximum bound</u> 15.0 Ma

<u>Suggested prior distribution</u> uniform

<u>Age justifications</u> There are no radiometric dates for the SHU 2-2 locality, and so its age estimate is based on geochronological comparisons with Asian and African faunas. These faunal correlations indicate an age between 8.0 and 6.5 Ma, with the most probable age reported as being around 7.0 Ma (Gilbert et al., 2014), but we prefer to use the minimum of this age range as our minimum bound here. As already discussed (see “Crown Cercopithecidae” and “Crown Colobinae” above), a diverse range of stem cercopithecoids, but no crown forms, are known from the early Miocene prior to ∼15 Ma (Locke et al., 2020: table 1), with the African fossil record becoming scarce 15-6 Ma (Rossie et al., 2013). A few fossils are known within this interval that may be relevant for calibrating this node, in particular a possible stem papionin from the Beticha locality of the Chorora Formation at 8.33-7.5 Ma (Suwa et al., 2015; Katoh et al., 2016). However, material of this Beticha taxon is extremely fragmentary (Suwa et al., 2015), it has (not to our knowledge) been included in a formal phylogenetic analysis, and Roos et al. (2019, p. 119) pointed out the difficulty in determining whether it is a stem papionin or stem cercopithecine without lower incisors that might reveal whether or not enamel was present lingually, (absence of lingual enamel is the only compelling dental synapomorphy of Papionini). For this reason, we do not use the Beticha taxon to provide our minimum bound.

We use 15.0 Ma as our maximum bound, based on the same reasoning as for crown Colobinae (see “Crown Colobinae” above). Because of the poor fossil record 15-6 Ma, and the possibility of a markedly earlier divergence (based on the Beticha taxon) than specified by our minimum bound, a uniform prior on this calibration seems most appropriate.

<u>Additional CladeAge calibration</u> AUH 1321 is the oldest known cercopithecin. Discounting the possible stem papionin from Beticha for the reasons discussed above, we consider the oldest record of Papionini to be two isolated teeth from the Lukeino Formation at Tugen Hills identified as cf. “*Parapapio*” *lothagamensis* (Gilbert et al., 2010). Although yet to be rigorously tested by a suitably comprehensive phylogenetic analysis, it is generally accepted that “*Parapapio*” *lothagamensis* is a stem papionin (Leakey et al., 2003; Pugh and Gilbert, 2018). The specimens from the Lukeino Formation described by Gilbert et al. (2010) closely resemble “*Parapapio*” *lothagamensis* in dimensions and overall morphology, and were identified by those authors as probably belonging to that taxon, or a slightly smaller close relative. The older of the two specimens (KNM-LU 861) comes from the Cheboit locality BPRP#29, which is ∼6.1 Ma (Gilbert et al., 2010), which we use as our CladeAge calibration.

<u>Comments</u> Our minimum bound on this node is the same as that proposed by Roos et al. (2019) in their calibration set 2, whilst dos Reis et al. (2018), used a younger minimum bound of 5.0 Ma, based on “*Parapapio” lothagamensis*; although the phylogenetic analyses of Gilbert et al. (2014) and Plavcan et al. (2019) differ somewhat, they both place AUH 1321 within crown Cercopithecini (see below), and so the Cercopithecini-Papionini split must predate this. For a maximum bound, Roos et al. (2019) used the ∼12.5 Ma Kabasero colobine material (Rossie et al., 2013; see “Crown Cercopithecidae” above), which, as discussed, we consider overly restrictive given the poor African primate record 15-6 Ma (Rossie et al., 2013). The maximum bound of dos Reis et al. (2018) meanwhile, was 23 Ma, based on the presence of *Kamoyapithecus* (which dos Reis et al., 2018, considered to be hominoid) at ∼25 Ma, and the appearance of the stem cercopithecid *Prohylobates* at 19.5 Ma onwards. Similarly to crown Colobinae (see above), we consider this overly conservative: the diversity of stem cercopithecoids but absence of crown forms in the early Miocene African record prior to ∼15 Ma persuades us that the Cercopithecini-Papionini split probably postdates this.

### 5.17 Crown Papionini = Macacina-Papionina split

Calibrating taxon cf. *Macaca* sp.

<u>Specimen</u> MGPT-PU 130508, a partial male cranium, from the Moncucco Torinese locality, Italy (Alba et al., 2014b).

<u>Phylogenetic justification</u> In a conference abstract, Alba et al. (2014a) reported that MGPT-PU 130508 is “undoubtedly papionin, as evidenced by facial and dental morphology and size”, and that its molars “display the typical generalized papionin morphology that is characteristic of *Macaca*, and their size fits with the upper-most range of *M. sylvanus* subspp.”, and they identified it as cf. *Macaca* sp. A full description of this significant specimen has yet to be published, and it lacks a full phylogenetic context, but based on the information provided by Alba et al. (2014a) we tentatively recognize this as a member of Macacina. In particular, we consider that it provides a more robust basis for calibrating this node than older (∼7.0-5.8 Ma) but much more fragmentary remains of ?*Macaca* sp. from Menacer, Algeria (Arambourg, 1959; Delson, 1975), which have been used by some previous authors (see below).

<u>Hard minimum bound</u> 5.33 Ma

<u>Soft maximum bound</u> 12.51 Ma

<u>Suggested prior distribution</u> uniform

<u>Age justifications</u> The fossil locality at Moncucco Torinese has been assigned a late Turolian (MN13, late Miocene) age based on its fossil fauna. The presence of an ostracod assemblage assigned to the *Loxocorniculina djafarovi* Zone allows a further refinement of the age to 5.40-5.33 Ma (Alba et al., 2014b), with the minimum age providing our minimum bound. For a maximum bound, we propose the maximum age of the oldest crown cercopithecid, namely the Kabasero Colobinae gen. et. sp. indet. material, which is 12.51 Ma (see “Crown Cercopithecidae” above); although the African primate fossil record is poor 15-6 Ma (Rossie et al., 2013), it seems unlikely that the Macacina-Papionina split, which is nested well within Cercopithecidae, would predate the overall oldest crown cercopithecid record.

We do not use the 7.0-5.8 Ma record of ?*Macaca* sp. from Menacer, Algeria (Arambourg, 1959; Delson, 1975) to calibrate this node (see below), but this record raises the possibility that our minimum bound is relatively conservative; we therefore propose a uniform prior on this calibration.

<u>Additional CladeAge calibration</u> We consider MGPT-PU 130508 to be the oldest robust record of Macacina. Based on available evidence, we consider the oldest robust record of Papionina to be 4.2-4.1 Ma old specimens from Kanapoi, West Turkana, Kenya, which have been identified as *Theropithecus* sp. indet. (Frost et al., 2020), and which provide our additional CladeAge calibration.

<u>Comments</u> dos Reis et al. (2018) did not calibrate this node. However, Roos et al. (2019) used an older minimum bound on this node of 5.8 Ma in their “calibration set 1” based on ∼7.0-5.8 Ma old remains of *?Macaca* sp. from Menacer, Algeria (Arambourg, 1959; Delson, 1975). Roos et al. (2019) noted themselves that it is unclear whether ?*Macaca* sp. from Menacer falls on the Macacina or the Papionina lineage. More seriously, Jablonski and Frost (2010) observed that there are no features of the ?*Macaca* sp. material from Menacer that would distinguish it from being a stem papionin, as was also noted by Delson (1975, 1980) and Szalay and Delson (1979). We therefore refrain from using this taxon for calibrating this node and instead use the slightly younger cf. *Macaca* from Moncucco Torinese discussed above. Roos et al. (2019) also used a comparatively young maximum bound of 8 Ma based on the possible stem papionin from the Beticha locality; we have already discussed the uncertainty surrounding this material (see “Crown Cercopithecinae” above), and such a tight maximum bound seems unjustified given the comparatively poor record of primates in Africa between 15 and 6 Ma (Rossie et al., 2013).

### 5.18 Crown Cercopithecini

<u>Calibrating taxon</u> Cercopithecini sp. indet.

<u>Specimen</u> AUH 1321, a lower left molar (most likely the first molar), from the SHU 2-2 locality in the Baynunah Formation, Abu Dhabi (Gilbert et al., 2014).

<u>Phylogenetic justification</u> As already mentioned (see “Crown Cercopithecinae” above), published phylogenetic analyses indicate that AUH 1321 is a crown cercopithecin: it was placed as sister to *Chlorocebus* or *Cercopithecus* in the analysis of Gilbert et al. (2014), but sister to

*Miopithecus* or in a polytomous clade with all extant cercopithecin genera except *Allenopithecus* in the analysis of Plavcan et al. (2019). A combination of features makes AUH 1321 most similar to non-*Allenopithecus* cercopithecins, namely a small and narrow molar with low-to-moderately flaring, elongated basin, and a distally expanded lophid (Gilbert et al., 2014). Nevertheless, the variation in the position of AUH 1321 between these analyses mean that its precise affinities are unclear. Furthermore, molecular phylogenies support a somewhat different set of relationships within Cercopithecini than do the analyses of Plavcan et al. (2019), in which the deepest split among extant cercopithecins is between *Allenopithecus* and the remaining genera; for example, Perelman et al. (2011) found *Allenopithecus* to be part of a clade that also includes *Chlorocebus* and *Erythrocebus*, whilst dos Reis et al. (2018) recovered an *Allenopithecus*+*Miopithecus* clade. These issues notwithstanding, we consider the phylogenetic analyses of Gilbert et al. (2014) and Plavcan et al. (2019) to collectively comprise sufficient evidence that AUH 1321 postdates the deepest split within Cercopithecini, and so can be used to provide a minimum bound on this node.

<u>Hard minimum bound</u> 6.5 Ma

<u>Soft maximum bound</u> 12.51 Ma

<u>Suggested prior distribution</u> uniform

<u>Age justifications</u> The age of the SHU 2-2 locality, which informs the minimum bound of this node, is discussed above (see “Crown Cercopithecinae”). Our maximum bound and suggested prior distribution follow the same logic as for crown Papionini (see above).

<u>Additional CladeAge calibration</u> Because it is uncertain exactly where AUH 1321 fits within crown Cercopithecini, and because of the incongruence between morphological (Gilbert et al., 2014; Plavcan et al., 2019) and molecular (e.g., Perelman et al., 2011; dos Reis et al., 2018) phylogenies of Cercopithecini, we refrain from suggesting an additional CladeAge calibration for this node.

<u>Comments</u> This node was not calibrated by Benton et al. (2015), dos Reis et al. (2018), or Roos et al. (2019).

### 5.19 Crown Hominoidea = Hominidae–Hylobatidae split

<u>Calibrating taxon</u> Kenyapithecus wickeri

<u>Specimen</u> KNM-FT 46a-b (holotype), left maxillary fragment with C1 and P4-M2 present, from Fort Ternan, Kenya (Leakey, 1961).

<u>Phylogenetic justification</u> *Kenyapithecus* has consistently been referred to as a crown hominoid, and specifically a hominid, by researchers (Pickford, 1985; Kelley et al., 2008; Harrison, 2010b; Alba, 2012) based in particular on the presence of the putative hominid synapomorphy of an anteriorly situated zygomatic root that is relatively high above the alveolar plane. This has been supported by formal phylogenetic analyses, with *Kenyapithecus* typically recovered as a stem hominid (e.g., Young and MacLatchy, 2004; Begun et al., 2012; Worthington, 2012), although Nengo et al. (2017) and Gilbert et al. (2020) found it to fall within crown Hominidae as a pongine. Regardless of its exact affinities, a position within crown Hominoidea for *Kenyapithecus* is well supported (but see Benoit and Thackeray, 2017), and we consider it the oldest definitive crown hominoid currently known.

<u>Hard minimum bound</u> 13.4 Ma

<u>Soft maximum bound</u> 25.235 Ma

<u>Suggested prior distribution</u> offset exponential

<u>Age justifications</u> The minimum age is based on dates of the Fort Ternan fossil locality published by Pickford et al. (2006a), who report on whole-rock K/Ar and single-crystal 40Ar/39Ar dates of lava flows underlying and overlying the fossil beds at Fort Ternan. The fossil beds at Fort Ternan are estimated to be 13.7 ± 0.3 Ma (Pickford et al., 2006a), giving an age range of 13.4-14.0 Ma, with the minimum age as our hard minimum bound. A second species of *Kenyapithecus, K. kizili*, has been described from Paşalar, Turkey (Kelley et al., 2008), which may be slightly older than *K. wickeri* (Roos et al., 2019). However, the age of Paşalar is poorly constrained (Casanovas-Vilar et al., 2011; Roos et al., 2019) and we do not use it to inform our minimum bound. The maximum bound is based on the maximum age of the oldest stem-hominoid *Rukwapithecus* (see “Crown Catarrhini” and “Crown Cercopithecidae” above).

In a situation equivalent to that seen in cercopithecoids (see “Crown Cercopithecidae”), the early Miocene African fossil record of Hominoidea is characterized by a diversity of stem taxa (proconsuline and nyanzapithecine “proconsulids”) without any evidence of crown representatives (Harrison, 2010b; Stevens et al., 2013; Nengo et al., 2017; Almécija et al., 2021). We tentatively interpret this as evidence that the Hominidae-Hylobatidae split was probably much closer to our minimum bound than our maximum bound, and so we propose calibrating this divergence with an offset exponential prior distribution. Assuming a 5% probability of exceeding the soft maximum bound, this would give a mean and median prior on this divergence of 17.4 and 16.1 Ma respectively.

<u>Additional CladeAge calibration</u> We consider *Kenyapithecus wickeri* to be the oldest known hominid (stem or crown, see above). The oldest hylobatid is the recently described *Kapi ramnagarensis* from the Lower Siwaliks of Ramnagar, India, with an estimated age of ∼13.8-12.5 Ma (Gilbert et al., 2020), which provides the additional CladeAge calibration for this node.

<u>Comments</u> Our minimum and maximum bounds are broadly similar to those of Roos et al. (2019). By contrast, Benton et al. (2015) proposed the crown hominid (stem pongine) *Sivapithecus* as the oldest crown hominoid, with a minimum age of 11.6 Ma, and used the age of the earliest anthropoids in the Fayum Depression as their maximum bound; in light of our discussion above, we consider both minimum and maximum bounds proposed for this node by Benton et al. (2015) to be unduly conservative. dos Reis et al. (2018) did not calibrate this node, but they stated in three separate places that they considered the ∼25 Ma old *Kamoyapithecus* to be a “crown hominoid”, a conclusion that they themselves admitted is “controversial”. However, the paper they cited in support of this conclusion, Zalmout et al. (2010), found *Kamoyapithecus* to be a stem (not crown) hominoid, and more distantly related to Hylobatidae+Hominidae than is “Proconsulidae”; table 1 of dos Reis et al. (2018) also lists *Kamoyapithecus* as a stem hominoid, in agreement with current evidence, as summarized above.

### 5.20 Crown Hominidae = Homininae-Ponginae split

<u>Calibrating taxon</u> Sivapithecus indicus

<u>Specimen</u> (GSP) Y 16075, a partial maxilla (Raza et al., 1983; Kappelman et al., 1991) with the connection between the maxilla and premaxilla partially preserved (Begun, 2015), from locality Y494 from the Chinji Formation, Pakistan (Pilgrim, 1910).

<u>Phylogenetic justification</u> *Sivapithecus* has been consistently recovered as a pongine in recent phylogenetic analyses (e.g., Begun et al., 2012; Nengo et al., 2017; Gilbert et al., 2020). (GSP) Y 16075 is the oldest *Sivapithecus* specimen to preserve the derived subnasal anatomy characteristic of modern orangutans (*Pongo* spp.; Kappelman et al., 1991). Isolated teeth from slightly older sites in the Chinji Formation have been referred to *Sivapithecus*, but they lack diagnostic features to support this referral (Kappelman et al., 1991), and so we do not use these for calibration purposes. We note that the slightly older *Kenyapithecus* (see “Crown Hominoidea” above) has been recovered as a pongine in some recent phylogenetic analyses (Nengo et al., 2017; Gilbert et al., 2020), but others place it as a stem-hominid (e.g., Young and MacLatchy, 2004; Begun et al., 2012; Worthington, 2012), and so it is not suitable for calibrating this node.

<u>Hard minimum age</u> 12.3 Ma

<u>Soft maximum age</u> 25.235 Ma

<u>Suggested prior distribution</u> offset exponential

<u>Age justifications</u> The minimum bound is based on the reported age of 12.3 Ma for another site in the mid-Chinji Formation, Y647 (which also preserves *Sivapithecus indicus* specimens), which is stated to be at the same stratigraphic level as Y494 (Morgan et al., 2015); this age is stated to be based on magnetostratigraphy, but Morgan et al. (2015) do not provide further details, and so it should be treated as tentative. The maximum bound is based on the maximum age of the oldest stem-hominoid *Rukwapithecus* (see “Crown Catarrhini”, “Crown Cercopithecidae”, and “Crown Hominoidea” above).We consider that this calibration is best modelled as an offset exponential distribution, based on the same arguments given for the crown Hominoidea node (see “Crown Hominoidea” above). Assuming a 5% probability of exceeding the soft maximum bound, this would give a mean and median prior on this divergence of 16.6 and 15.3 Ma respectively.

<u>Additional CladeAge calibration</u> We consider *Sivapithecus indicus* to be the oldest definitive pongine. A number of “dryopiths” (sensu Almécija et al., 2021; Urciuoli et al., 2021) from Europe, including the ∼9.6-8.7 Ma old *Ouranopithecus macedoniensis* (Sen et al., 2000; Koufos et al., 2016), have been found to be stem hominines in some published phylogenetic analyses (e.g., Begun et al., 2012), but not others (e.g., Alba et al., 2015), and their position appears sensitive to analytical assumptions (Young and MacLatchy, 2004; Worthington, 2012). Given the ongoing controversy regarding the affinities of the European “dryopiths” (see Benoit and Thackeray, 2017; Fuss et al., 2018; Almécija et al., 2021), we follow Gilbert et al. (2020) in viewing them as Hominoidea incertae sedis, and do not use them as the oldest record of Homininae. Instead, we use the slightly younger *Chororapithecus abyssinicus*, which we recognize as a crown hominine (stem gorillin), with an age range of 8.33-7.5 Ma (see “Crown Homininae” below).

<u>Comments</u> Roos et al. (2019) used *Kenyapithecus wickeri* (with a maximum age of 14.9 Ma) as their maximum bound on this node, but given that Nengo et al. (2017) and Gilbert et al. (2020) found *Kenyapithecus* to be a pongine, this may be overly restrictive, and we take a more conservative approach, instead using the maximum age of the oldest stem-hominoid *Rukwapithecus fleaglei* for setting the soft maximum bound. Benton et al. (2015), meanwhile, used a maximum of 33.9 Ma based on the age of the oldest known crown anthropoids from the L-41 Quarry of the Fayum, Egypt (see “Crown Anthropoidea” above), which seems excessively conservative given the diversity of stem hominoids but absence of crown forms in the early Miocene African record (see “Crown Hominoidea” above).

### 5.21 Crown Homininae = Gorillini-Hominini split

<u>Calibrating taxon</u> Chororapithecus abyssinicus

<u>Specimen</u> CHO-BT 4 (holotype), a right M2, from the Chorora Formation at Beticha, Ethiopia (Suwa et al., 2007).

<u>Phylogenetic justification</u> Although *Chororapithecus* has not, to our knowledge, been included in a formal phylogenetic analysis, Suwa et al. (2007) discussed a number of apomorphic dental features that are shared by *Chororapithecus* and *Gorilla* (albeit that are less derived in the fossil taxon), and they noted that no other fossil hominoid besides the enigmatic *Oreopithecus* from the Miocene of Italy exhibits the same combination of apomorphies. Some subsequent authors (e.g., Begun et al., 2012; Fuss et al., 2018) have concluded that the relationship between *Chororapithecus* and *Gorilla* is questionable based on the limited fossil evidence, and Suwa et al. (2007) themselves noted that the derived resemblances may be homoplastic. However, in the absence of specific evidence to the contrary (such as a formal phylogenetic analysis that includes *Chororapithecus*, *Gorilla*, and a suitably dense sampling of other crown and stem hominoids and appropriate outgroups), we tentatively accept that C. *abyssinicus* is a stem gorillin, following Suwa et al. (2007).

<u>Hard minimum bound</u> 7.5 Ma

<u>Soft maximum bound 15</u> Ma

<u>Suggested prior distribution</u> uniform

<u>Age justifications</u> In their original description of *Chororapithecus abyssinicus*, Suwa et al. (2007) identified the Beticha locality as being 10.5-10.1 Ma old. Subsequent radiometric dating now indicates a maximum of 8.33 Ma and a minimum of 7.5 Ma for this locality (Katoh et al., 2016; see “Crown Cercopithecinae” above), and we use the latter date as our minimum bound. For our maximum bound, we use the same 15 Ma date as for crown Colobinae and crown Cercopithecinae (see above), reflecting the generally poor record of primates in Africa 15-6 Ma (Rossie et al., 2013), and the fact that stem hominoids were diverse but crown hominoids were apparently absent in Africa during the early Miocene prior to 15 Ma.

Although we do not use this specimen for calibration purposes, Pickford and Senut’s (2005) report of a ∼12.5 Ma isolated lower molar from the Ngorora Formation that they suggest may belong to the *Pan* lineage (but which Kunimatsu et al. (2007) considered resembles *Gorilla*) raises the possibility that this divergence may be markedly older than our minimum bound. Katoh et al. (2016), meanwhile, suggested that the ∼9.8 Ma old *Nakalipithecus nakayamai* may be directly ancestral to *Chororapithecus*; given the limited evidence in support of this relationship provided by Katoh et al. (2016), we have not used *Nakalipithecus* to calibrate this node, but we recognize that it may indicate a markedly earlier time for this divergence than specified by our minimum bound. Considering these issues, and again taking into account the generally poor African primate record 15-6 Ma (Rossie et al., 2013), we consider that this calibration is most appropriately modelled as a uniform prior distribution.

<u>Additional CladeAge calibration</u> We recognize *Chororapithecus abyssinicus* as the oldest gorillin. The oldest member of the *Homo* lineage that is supported by robust dating and phylogenetic analyses is *Ardipithecus ramidus* (see “*Homo-Pan* split” below). The oldest well-dated *A. ramidus* material is 4.799-4.631 Ma old (Semaw et al., 2005; Simpson et al., 2019; see “Homo lineage-Pan lineage split” below), and provides our additional CladeAge calibration for this node.

<u>Comments</u> Benton et al. (2015) did not calibrate this node, but both dos Reis et al. (2018) and Roos et al. (2019) used a similar minimum bound to that proposed here, based on *Chororapithecus*. However, both dos Reis et al. (2018) and Roos et al. (2019) based their maximum bound on the age of *Sivapithecus*, and we argue that *Sivapithecus* and other Eurasian hominoids do not provide a clear basis for constraining the crown Homininae or crown Hominini nodes, both of which probably took place in Africa, hence our decision to use the African fossil record to inform our maximum bound (see “*Homo*-*Pan* split” below for more detailed discussion).

### 5.22 Homo*-*Pan *split*

<u>Calibrating taxon</u> Ardipithecus ramidus

<u>Specimen</u> WMN5sw/P56, a mandibular ramus and partial dentition (p3-m3) from GMW5sw locality in Gona, Ethiopia (Semaw et al., 2005; Simpson et al., 2019).

<u>Phylogenetic justification</u> Although *Ardipithecus ramidus* preserves numerous similarities to the chimpanzee condition in cranial, dental and postcranial material, White et al. (1995) argued that some, if not all, of these are primitive retentions from the last common ancestor of chimpanzees and humans. Notable features of *A. ramidus* that appear to be synapomorphies placing it as a member of the *Homo* lineage include the more incisiform canines, an anteriorly located foramen magnum, and a proximal ulnar morphology that is shared with *Australopithecus* species (White et al., 1995, 2009; Suwa et al., 2009a, 2009b; but see Harrison, 2010c). This interpretation has been tested in phylogenetic analyses by Dembo et al. (2015, 2016), Mongle et al. (2019), and Püschel et al. (2021), who all recovered *A. ramidus* as sister taxon to all later members of the *Homo* lineage. Although *Ardipithecus kadabba* is slightly older than *A. ramidus* (5.8-5.2 Ma, Haile-Selassie, 2001; WoldeGabriel et al., 2001; 2004), we refrain from using this species to calibrate this node as it has not been included in any of these phylogenetic analyses, most likely due to the scarcity of *Ardipithecus kadabba* material. The phylogenetic analyses of Dembo et al. (2015, 2016), Mongle et al. (2019), and Püschel et al. (2021) all placed *Sahelanthropus* closer to *Homo* than to *Pan*, but doubts over the stratigraphic provenance of *Sahelanthropus*, and hence its age, mean that we do not use it as our calibrating taxon here (see “Comments” below).

<u>Hard minimum bound</u> 4.631 Ma

<u>Soft maximum bound</u> 15 Ma

<u>Suggested prior distribution</u> uniform

<u>Age justifications</u> The oldest *A. ramidus* localities (GMW1, GMW5sw, and GWM9) have been assigned to the C3n.2r magnetozone (Simpson et al., 2019) which corresponds to an age of 4.799-4.631 Ma (Hilgen et al., 2012). We have already discussed the relevance of the poor record of fossil primates in Africa 15-6 Ma (Rossie et al., 2013), and the ∼12.5 Ma old Ngorora tooth that Pickford and Senut (2005) suggested might be a member of the *Pan* lineage. Based on these factors, we again prefer to take a conservative approach for this node and suggest the same maximum bound as for Homininae (see “Crown Homininae” above). For the same reason, we also suggest a uniform bound is the appropriate prior distribution for this node.

<u>Additional CladeAge calibration</u> We recognize *Ardipithecus ramidus* as the oldest known member of the *Homo* lineage that has a well-constrained age. The fossil record of its sister-clade, the *Pan* lineage, is extremely limited. To date, the oldest fossils are specimens that have been referred to the modern genus *Pan* from the Kapthurin Formation of Kenya (although this was questioned by Harrison, 2010b, who instead argued that they may belong to *Homo*), the age of is constrained by 40Ar/39Ar dates of 545 ± 3 kyr for deposits underlying the fossils and 284 ± 12 kyr for deposits overlying them (Deino and McBrearty, 2002; McBrearty and Jablonski, 2005). The fossils are located most closely to the underlying deposit, and McBrearty and Jablonski (2005) argued that their age is likely to be close to 0.5 Ma. However, we prefer to use the entire age range (including confidence intervals) for this record, giving an additional CladeAge calibration for this node of 0.548-0.272 Ma.

<u>Comments</u> Unlike us, Benton et al. (2015), dos Reis et al. (2018), and Roos et al. (2019) all used the age of *Sahelanthropus* to provide a minimum bound of 6.2 Ma (Roos et al., 2019), 6.5 Ma (Benton et al., 2015), or 7.5 Ma (dos Reis et al., 2018) on this divergence. *Sahelanthropus* has various synapomorphies in its cranial morphology placing it closer to the *Homo* lineage than the *Pan* lineage (see e.g., Brunet et al., 2002; Zollikofer et al., 2005; MacLatchy et al., 2010; Emonet et al., 2014). This has been questioned by some authors (Wolpoff et al., 2002; Wolpoff and Pickford, 2006), but recent phylogenetic analyses by Dembo et al. (2015, 2016), Mongle et al. (2019), and Püschel et al. (2021) have consistently placed *Sahelanthropus* closer to *Homo* than to *Pan*, as the basalmost member of the *Homo* lineage among the taxa included in these analyses. However, questions have been raised about the stratigraphic origin of *Sahelanthropus* material (Beauvilain, 2008), and thereby on its reported age of 7.2-6.8 Ma (Lebatard et al., 2008). Ahern (2018) concluded that the fossil material of *Sahelanthropus* is most likely of late Miocene age, but suggested that its age could not be constrained more accurately than 7.5-5.0 Ma based on available data. We accept that *Sahelanthropus* is most likely a member of the *Homo* lineage, but due to the uncertainty surrounding its age we prefer to take a more conservative approach and use the more securely dated *Ardipithecus ramidus* for calibrating this node. However, if more securely constrained *Sahelanthropus* material is reported that predates the age of *A. ramidus*, the minimum age of this calibration will need to be updated.

Benton et al. (2015) proposed a maximum bound of 10 Ma on this node, given that “a range of ape taxa, *Ankarapithecus* from Turkey (10 Ma), *Gigantopithecus* from China (8–0.3 Ma), *Lufengopithecus* from China (10 Ma), *Ouranopithecus* from Greece (10–9 Ma), and *Sivapithecus* from Pakistan (10–7 Ma) give maximum ages of 10 Ma, early in the late Miocene, and these deposits have yielded no fossils attributable to either chimps or humans”. Importantly, however, all of these taxa are Eurasian not African, and current evidence supports relatively limited dispersal of hominoids out of Africa (Gilbert et al., 2020). In particular, it seems likely that the split between the *Homo* and *Pan* lineages occurred in Africa (but see Fuss et al., 2018), with members of the *Homo* lineage probably not dispersing out of Africa until the early Pleistocene (Trifonov et al., 2019), and members of the *Pan* lineage apparently never doing so. Thus, the absence of members of the *Homo* and *Pan* lineages in ∼10 Ma old Eurasian deposits would still be expected even if the split between these lineages had already occurred by this time, and so there seems no compelling reason to use this as a maximum bound on this node. Roos et al. (2019), meanwhile, assumed a maximum bound of 8 Ma based on the stem-gorillin *Chororapithecus*, and dos Reis et al. (2018) also used *Chororapithecus* to provide a maximum bound on this divergence, but assumed a 10 Ma age for *Chororapithecus* based on the initial report by Suwa et al. (2007), whereas its age has now been revised down to 7.5-8.33 Ma (Katoh et al., 2016). Regardless, we consider use of *Chororapithecus* to inform the maximum bound on this node to be excessively restrictive, given the ∼12.5 Ma potential *Pan* relative described by Pickford and Senut (2005), and the overall poor African record of primates 15-6 Ma (Rossie et al., 2013), discussed above.

### 5.23 Crown Platyrrhini = Pitheciidae-(Aotidae+Atelidae+Callitrichidae+Cebidae) split

<u>Calibrating taxon</u> Stirtonia victoriae

<u>Specimen</u> Duke/INGEOMINAS 85-400 (holotype), a right maxilla preserving erupted dP2-dP4 M1-M2, and mineralized but unerupted C and P2-P4, from Duke Locality 28, La Venta, Colombia (Kay et al., 1987).

<u>Phylogenetic justification</u> In our tip-dated total evidence phylogenetic analyses, the oldest well supported member of crown Platyrrhini is *Stirtonia*, which was strongly supported as sister to *Alouatta*, within crown Atelidae (Figures 1-2). The only potentially older taxon placed within crown Platyrrhini in our analyses was the 19.76-10.4 Ma old *Proteropithecia neuquenensis* from the Collón Curá Formation of Argentina (Kay et al., 1998; Kay and Perry, 2019, see Supplementary Online Material for discussion of the age of this specimen). However, support for placing *Proteropithecia* within crown Platyrrhini was very low (BPP = 0.30), and in fact there was greater support for placing it outside the crown clade (see “Results” and “Discussion” above). All older fossil platyrrhines, including *Panamacebus transitus* (which has been found to be a cebid in some other published analyses; (Bloch et al., 2016; Kay et al., 2019), were placed outside the crown clade (see “Results” and “Discussion” above).

<u>Hard minimum bound</u> 13.363 Ma

<u>Soft maximum bound</u> none

<u>Suggested prior distribution</u> not applicable (minimum bound only)

<u>Age justifications</u> *Stirtonia victoriae* is currently the oldest *Stirtonia* species known, with all known material from Duke Locality 28 at La Venta, approximately 290m below the stratigraphic level from where specimens of the younger *S. tatacoensis* have been collected (Kay et al., 1987). According to Guerrero (1997, fig. 2.8), Duke Locality 28 is within the Cerro Gordo Beds of the La Victoria Formation. Montes et al. (2021, fig. 3) indicated that the Cerro Gordo Beds, the overlying Chunchullo Beds, and the underlying San Alfonso Beds all lie within Chron C5ABn, which spans from 13.608 to 13.363 Ma (Gradstein et al., 2012), with the latter providing our minimum bound.

For less inclusive divergences within crown Platyrrhini (primarily divergences within families; see the calibrations that follow), we have proposed a maximum bound based on the maximum reported age of the oldest probable stem platyrrhine specimens, which is 34.5 Ma (Antoine et al., 2021), although we note that this date has been questioned (Campbell et al., 2021). The ancestor of crown Platyrrhini was probably a very small (∼400g), insectivore-frugivore (Lynch Alfaro, 2017; Silvestro et al., 2019b) that morphologically is likely to have been little different from the specimens described by Antoine et al. (2021). This, together with the overall poor record of platyrrhines, means that it is difficult to rule out an early (pre-Oligocene) origin for crown Platyrrhini on fossil grounds alone; for this reason, we do not propose a maximum bound for this calibration.

<u>Additional CladeAge calibration</u> *Stirtonia victoriae* is the oldest known atelid (and is, in fact, a crown form, closer to the alouattine *Alouatta* than to atelines), and so is the oldest known member of the Aotidae+Atelidae+Callitrichidae+Cebidae clade. Ignoring *Proteropithecia* for the reasons already discussed (see “Results and “Discussion” above), the oldest known member of its sister clade, Pitheciidae, is the crown pitheciid *Cebupithecia sarmientoi*, which is 13.183-13.032 Ma old (see “Crown Pitheciidae” below).

<u>Comments</u> Benton et al. (2015) did not calibrate this node, but dos Reis et al. (2018) proposed a minimum of 15.7 Ma based on the age of *Proteropithecia* reported by Kay et al. (1998); as noted above, our total evidence tip-dating analyses do not clearly support *Proteropithecia* as a crown platyrrhine. dos Reis et al.’s (2018) maximum bound was 33 Ma, based on the age of the oldest crown anthropoid, the stem catarrhine *Catopithecus* (see “Crown Anthropoidea” above).

However, if the 34.5 Ma date reported by Antoine et al. (2021) is correct, then the oldest record of probable platyrrhines predates this (although we accept that these are almost certainly stem forms). Overall, we consider the platyrrhine fossil record to be too incomplete to confidently apply a maximum bound on this node.

### 5.24 Crown Pitheciidae = Callicebinae-Pitheciinae split

<u>Calibrating taxon</u> Cebupithecia sarmientoi

<u>Specimen</u> UCMP 38762 (holotype), a nearly complete skull, mandible, axial skeleton, and limb bones, from the Monkey Beds at La Venta, Colombia (Stirton and Savage, 1951).

<u>Phylogenetic justification</u> Numerous synapomorphies support *Cebupithecia* as a pitheciine (Stirton and Savage, 1951; Orlosky, 1974; Rosenberger, 1979; Kay, 1990), and it has been consistently placed within crown Pitheciidae as a stem pitheciine in published phylogenetic analyses (Kay, 2015; Bloch et al., 2016; Marivaux et al., 2016; Kay et al., 2019; Ni et al., 2019: fig. S1). Our own tip-dating analyses also support *Cebupithecia* as a stem-pitheciine, with relatively strong support (see “Results” and “Discussion” above). A second fossil platyrrhine from La Venta, *Nuciruptor rubricae*, has also been consistently placed as a stem pitheciine in published phylogenetic analyses (Kay, 2015; Bloch et al., 2016; Marivaux et al., 2016; Kay et al., 2019), including the mandibular analysis presented here. However, material of *Nuciruptor* is from the El Cardon Red Beds, which are younger than the Monkey Beds (12.829-12.272 Ma; Montes et al., 2021). A potentially older taxon, 19.76-10.4 Ma old *Proteropithecia neuquenensis* from the Collón Curá Formation of Argentina (Kay et al., 1998; Kay and Perry, 2019, see Supplementary Online Material for discussion of the age of this specimen) was found to be a stem pitheciine in recent phylogenetic analyses (Kay, 2015; Bloch et al., 2016; Marivaux et al., 2016; Kay et al., 2019). However, as noted above (see “Results” and “Discussion”), our mandibular analysis did not robustly support pitheciine affinities for *Proteropithecia*, and in any case the minimum age for this taxon is younger than for *Cebupithecia* (see Supplementary Online Material). Thus, we have not used *Proteropithecia* to calibrate this node.

<u>Hard minimum bound</u> 13.032 Ma

<u>Soft maximum bound</u> 34.5 Ma

<u>Suggested prior distribution</u> uniform

<u>Age justifications</u> The type specimen of *Cebupithecia sarmientoi* comes from the Monkey Beds at La Venta that correspond to the normal interval of Chron C5AA (Flynn et al., 1996; Kay and Madden, 1997). This interval spans from 13.183 to 13.032 Ma (Gradstein et al., 2012), with the latter date providing our hard minimum bound.

Our total evidence tip-dating analyses suggest that the most recent common ancestor of crown Platyrrhini is ∼21-28 Ma old, but the oldest known crown platyrrhines (including *Cebupithecia*) are some 10-15 Ma younger; all fossil platyrrhines older than ∼14 Ma are placed outside the crown in our analyses (see “Results” and “Discussion” above). Thus, the early stages of the evolution of crown Platyrrhini appear to be currently unsampled, probably because they occurred in northern South America, where the fossil record for this time period remains poor (although ongoing research is starting to improve this; e.g., Antoine et al., 2012, 2017; Bond et al., 2015; Bloch et al., 2016; Marivaux et al., 2016; Kay et al., 2019). For this reason, we suggest a conservative maximum bound of 34.5 Ma, based on the maximum reported age of the oldest platyrrhine specimens from TAR-21 site, Shapaja, Peru (Antoine et al., 2021); these specimens appear to be highly plesiomorphic, and similar to the better preserved *Perupithecus* (Antoine et al., 2021), which has been dated to 29.6 ± 0.08 Ma *(Campbell et al., 2021)*. Campbell et al. (2021) questioned the late Eocene age for TAR-21, and presented tip-dating analyses of fossil rodents suggesting an Oligocene age for this and other Shapaja sites. However, pending further exploration of this issue, we prefer to use the older age for TAR-21 as the maximum bound of this calibration as a conservative maximum. Given the obvious incompleteness of the fossil record, we also suggest that this should be modelled as a uniform distribution. We propose the same maximum bound and uniform prior distribution for all other divergences within Platyrrhini.

<u>Additional CladeAge calibration</u> We recognize *Cebupithecia sarmientoi* as the oldest stem pitheciine. The fossil taxon *Miocallicebus villaviejai* has been described as being dentally similar to, but much larger than, *Callicebus sensu lato* (= the currently recognized modern callicebine genera *Callicebus*, *Plecturocebus* and *Cheracebus*; (Takai et al., 2001; Kay, 2015). The only known specimen (IGM-KU 97001) is a partial maxilla preserving only a single fully intact tooth (M2), which is heavily worn, and its affinities have not been tested via formal phylogenetic analysis. Nevertheless, we consider the available evidence sufficient to recognize *Miocallicebus* as a fossil callicebine, and so we propose it as an additional CladeAge calibration here. IGM-KU 97001 comes from the Bolivia Site at La Venta, which is just above the Tatacoa Beds, towards the top of the La Victoria Formation. According to Montes et al. (2021, fig. 3), the base of the Tatacoa Beds is within Chron C5ABr, whilst the top of the La Victoria Formation is within Chron C5AAn. Following Gradstein et al. (2012), this gives an age range of 13.363-13.032 Ma for *Miocallicebus villaviejai*, which we use as our CladeAge calibration for the oldest record of Callicebinae.

### 5.25 Callitrichidae-Cebidae split

<u>Calibrating taxon</u> Lagonimico conclucatus

<u>Specimen</u> IGM 184531 (holotype), a crushed skull with partial upper dentition present and a near complete mandible with most of the mandibular dentition from Duke/INGEOMINAS locality 90 in the Victoria Formation, La Venta, Colombia (Kay, 1994).

<u>Phylogenetic justification</u> *Lagonimico* shares a number of dental synapomorphies with extant crown callitrichids (Kay, 1994: table 7), and recent phylogenetic analyses consistently place it as a stem-callitrichid (Kay, 2015; Bloch et al., 2016; Marivaux et al., 2016; Kay et al., 2019). We found a similar result in our tip-dating analyses, with *Lagonimico* strongly supported as sister to crown-callitrichids in both the mandibular and maxillary analysis, and so it is suitable for calibrating this divergence. Our analyses, as well as several others (Kay, 2015; Bloch et al., 2016; Marivaux et al., 2016; Kay et al., 2019) suggest that two other taxa from La Venta - *Mohanamico hershkovitzi* (Luchterhand et al., 1986) and *“Aotus” dindensis* (originally described as an aotid; Setoguchi and Rosenberger, 1987; see also Ni et al., 2019 fig. S1) - may also be stem callitrichids, but both are from the “Monkey Beds”, which are stratigraphically younger than Duke/INGEOMINAS locality 90 (Flynn et al., 1996; Montes et al., 2021; see below).

<u>Hard minimum bound</u> 13.183 Ma

<u>Soft maximum bound</u> 34.5 Ma

<u>Suggested prior distribution</u> uniform

<u>Age justifications</u> Duke/INGEOMINAS locality 90, source of IGM 184531, is located stratigraphically between the overlying Tatacoa Beds and underlying Chunchullo Beds of the Victoria Formation (Kay, 1994, Table 7; Montes et al., 2021). According to Montes et al. (2021, fig. 3), the base of the Tatacoa Beds is within Chron C5ABr, whilst the Chunchullo Beds are entirely within Chron C5ABn, as are the Cerro Gordo Beds that underlie them. Thus, Duke/INGEOMINAS Locality 90 must be younger than the base of Chron C5ABn, and older than the top of Chron C5ABr, which gives an age range of 13.608 to 13.183 Ma, with the latter providing our minimum bound.

Based on the poor record of crown Platyrrhini (see “Crown Pitheciidae” above), we suggest a conservative maximum bound of 34.5 Ma for this node, based on the maximum reported age of the oldest probable platyrrhine specimens described to date (Antoine et al., 2021), and a uniform prior distribution.

<u>Additional CladeAge calibration</u> *Lagonimico conclucatus* is the oldest known callitrichid. The sister taxon of Callitrichidae is either Cebidae, Aotidae, or Cebidae+Aotidae, as the precise relationships of *Aotus* have proven difficult to resolve (Osterholz et al., 2009; Perez et al., 2012; Valencia et al., 2018; Schrago and Seuánez, 2019; X. Wang et al., 2019; Vanderpool et al., 2020). Considering the fossil record of cebids and aotids together, the oldest well-supported representative of either family is *Neosaimiri*, which has been consistently proposed to be closely related to the modern *Saimiri* from its original description by Stirton (1951) onwards (e.g., Rosenberger et al., 1991; Takai, 1994). Congruent with this, recent phylogenetic analyses support *Neosaimiri* as a crown cebid, sister to *Saimiri* (Kay, 2015; Bloch et al., 2016; Marivaux et al., 2016; Kay et al., 2019), and we found the same relationship in our tip-dating analyses. The oldest *Neosaimiri* material, including the holotype UCMP 39205, comes from the Monkey Beds in the Villavieja Formation (see “Crown Cebidae” below), which 13.183 to 13.032 Ma (see “Crown Pitheciidae” above), and this age range provides our additional CladeAge calibration.

<u>Comments</u> Kay (2015) discussed records of two possible crown callitrichids from La Venta: an isolated upper incisor (IGM-KU 8402) and a lower fourth premolar (IGM-KU 8403) from the “Monkey Beds” that Setoguchi and Rosenberger (1985) tentatively referred to *Micodon kiotensis* (see also Rosenberger et al., 1990); and the holotype of *Patasola magdalenae* (IGM 184332), a partial lower jaw preserving the deciduous premolars and the molars from the stratigraphically older Duke Locality 40 (Kay and Meldrum, 1997), which is in the interval between the overlying Chunchullo Beds and underlying Cerro Gordo beds (and so appears to fall within Chron C5ABn; Montes et al., 2021). However, as discussed by Setoguchi and Rosenberger (1985), Rosenberger et al. (1990) and Kay and Meldrum (1997), the evidence for referring GM-KU 8402 and 8403 to Callitrichidae (whether stem or crown) is weak, and this proposed relationship has not been tested via formal phylogenetic analysis. By contrast, Kay and Meldrum (1997) did formally test the relationships of *Patasola* via maximum parsimony analysis of 55 dental characters and found that it fell within crown Callitrichidae. However, the overall topology for extant callitrichids recovered by Kay and Meldrum (1997) - namely (*Callimico*,(*Leontopithecus*,(*Saguinus*,*Callithrix*))) - is highly incongruent with molecular data (e.g., Garbino and Martins-Junior, 2018), which strongly supports (*Saguinus*,(*Leontopithecus*,(*Callimico*,*Callithrix*). Given this incongruence, which is likely to have a major impact on the polarities of the dental characters used by Kay and Meldrum (1997) to place *Patasola,* we do not consider the phylogenetic analysis of Kay and Meldrum (1997) to be strong evidence that *Patasola* is a crown callitrichid. We prefer not to propose calibrations for any divergences within crown Callitrichidae.

### 5.26 Crown Cebidae = Cebinae-Saimirinae split

<u>Calibrating taxon</u> *Neosaimiri fieldsi*

<u>Specimen</u> UCMP 39205 (holotype), comprising a left hemi-mandible preserving p2-m2, and a right hemi-mandible preserving i2-m2, from UCMP locality V4517 in the “Monkey Beds” of the Villavieja Formation at La Venta, Colombia (Stirton, 1951).

<u>Phylogenetic justification</u> As already discussed (see “Callitrichidae-Cebidae split” above), *Neosaimiri* has been consistently identified as a close relative of the extant saimirine genus *Saimiri* since its original description. Indeed, Rosenberger et al. (1991) concluded that *Neosaimiri* could be synonymized with *Saimiri*, although Takai (1994) argued that the two genera should be maintained as distinct. Regardless, a close relationship between *Neosaimiri* and *Saimiri* to the exclusion of *Cebus*, within crown Cebidae, has been a consistent feature of recent published phylogenetic analyses (Kay, 2015; Bloch et al., 2016; Marivaux et al., 2016; Kay et al., 2019). Our own tip-dating total evidence analyses find very strong support for a *Saimiri*+*Neosaimiri* clade to the exclusion of *Cebus* (see “Results” and “Discussion” above). The ∼20.9 Ma old *Panamacebus transitus* has also been placed within crown Cebidae in some published phylogenetic analyses (Bloch et al., 2016; Marivaux et al., 2016; Kay et al., 2019), but it fell outside crown Platyrrhini in our tip-dating analyses (see “Results” and “Discussion” above), and we do not use this taxon for calibrating purposes here.

<u>Hard minimum bound</u> 13.032 Ma

<u>Soft maximum bound</u> 34.5 Ma

<u>Suggested prior distribution</u> uniform

<u>Age justifications</u> As already discussed (see “Crown Pitheciidae” above), the age of the “Monkey Beds” at La Venta can be constrained to between 13.183 and 13.032 Ma. We propose the same maximum bound and uniform prior distribution as for crown Pitheciidae and the Callitrichidae-Cebidae split (see above).

<u>Additional CladeAge calibration</u> As discussed, we consider *Neosaimiri fieldsi* to be the oldest well-supported saimirine. If *Panamacebus* is discounted, the oldest well-supported member of Cebinae is *Acrecebus fraileyi* (Kay and Cozzuol, 2006) which has been consistently placed sister to *Cebus* in recent published phylogenetic analyses (Kay, 2015; Bloch et al., 2016; Marivaux et al., 2016; Kay et al., 2019). We recovered the same relationship in our tip-dating analyses. *Acrecebus fraileyi* is known from a specimen, LACM 134880 (a left M2) from locality LACM 5158 (“Bandeira”), Solimoes Formation, Acre River, Acre, Brazil. The age of the Acre vertebrate fauna of the Solimoes Formation has been controversial and remains poorly constrained (Cozzuol, 2006). The Patos locality, which is near to the Bandeira locality (Negri et al., 2010, fig. 15.1), has recently been proposed to be no older than 7 Ma based on palynological data (Leite et al., 2021). However, in the absence of more precise stratigraphic evidence, we follow Kay and Cozzuol (2006) and (2006; Tomassini et al., 2013) (2013) in assigning the Acre vertebrate fauna to the Huayquerian SALMA, which is 9.0-5.28 Ma following Prevosti and Forasiepi (2018), and provides our CladeAge calibration.

### 5.27 Crown Atelidae = Alouattinae-Atelinae split

<u>Calibrating taxon</u> Stirtonia victoriae

<u>Specimen</u> Duke/INGEOMINAS 85-400 (holotype), a right maxilla preserving erupted dP2-dP4 M1-M2, and mineralized but unerupted C and P2-P4, from Duke Locality 28, La Venta, Colombia (Kay et al., 1987).

<u>Phylogenetic justification</u> Numerous authors have noted that *Stirtonia* shares numerous dental similarities (at least some of them derived) with the modern genus *Alouatta* (Stirton, 1951; Rosenberger, 1979; Setoguchi et al., 1981; Kay and Cozzuol, 2006), and a *Stirtonia+Alouatta* clade has been recovered in several recent phylogenetic analyses (Kay and Cozzuol, 2006; Bloch et al., 2016; Marivaux et al., 2016; Kay et al., 2019). Our own total evidence tip-dating analyses also support this.

<u>Hard minimum bound</u> 13.363 Ma

<u>Soft maximum bound</u> 34.5 Ma

<u>Suggested prior distribution</u> uniform

<u>Age justifications</u> See “Crown Platyrrhini” above for discussion of the age of *Stirtonia victoriae*.

<u>Additional CladeAge calibration</u> *Stirtonia victoriae* is the oldest known alouattine. Kay and Cozzuol (2006) named *Solimoea acrensis* based on an isolated left m1 (the holotype, UFAC-LPP 5177) and an isolated right maxillary fragment preserving P3-4 (UFAC-LPP 5178) from the Patos locality (equivalent to LACM 4611) in the Solimoes Formation and identified it as an ateline. They also carried out a four different maximum parsimony analyses based of 57 dental characters (although only 25 of these were parsimony informative; Kay and Cozzuol, 2006: table 1) and using a molecular scaffold that was based on the studies of Meireles et al. (1999 b; Carla M. Meireles et al., 1999 a) but which is still in agreement with current molecular evidence (e.g. dos Reis et al., 2018): in all four analyses, *Solimoea* formed a clade with living atelines, with moderate-to-high (57-86%) bootstrap support depending on the analysis. Kay (2015) subsequently stated that he considered *Solimoea* to be specifically related to *Lagothrix* within crown Atelinae, although Kay and Cozzuol (2006) found that *Solimoea* fell outside crown Atelinae in three out of four of their phylogenetic analyses. Rosenberger et al. (2015) argued that *Solimoea* is more likely an alouattine, and cast doubt on whether the holotype m1 and the referred maxillary fragment represent the same taxon. However, in the absence of formal phylogenetic analysis supporting alternative relationships for *Solimoea*, we tentatively accept it as the oldest known ateline. Based on palynological evidence, the Patos locality is 7 Ma or younger (Leite et al., 2021); given that we accept a Huayquerian age for the Acre vertebrate fauna as a whole (see “Crown Cebidae” above) which is 9-5.28 Ma, we assign *Solimoea acrensis* an age range of 7-5.28 Ma for our additional CladeAge calibration.

## 6. Conclusion

Marjanović (2021) noted that compendia of fossil calibrations quickly go out of date, due both to the discovery of new fossils and to reinterpretation and reanalysis of those already known. However, the impact of this on analyses that need to use fossil calibration information can (we hope) be lessened by careful consideration of the appropriate prior distribution for each calibrated node, to adequately reflect our current uncertainty and to take into account the likely impact of future discoveries. For example, it is certainly possible, or even probable, that crown primates older than *Teilhardina brandti* will be discovered, but we think it highly unlikely that they will be found in the Cretaceous, an assumption that is taken into account by our suggested prior distribution on the age of crown Primates. Thus, we expect that improvements in the primate fossil record will lead to tighter constraints on the ages of particular nodes (mainly due to older minimum bounds), but not ones that actively conflict with those proposed here. In turn, revisions to this list should lead to more precise, but not contradictory, estimates of divergence times in future node-dating analyses.

## Acknowledgments

We thank the following people for discussion, for providing references, or both: David Alba, Pierre-Olivier Antoine, Ornella Bertrand, Jean Boubli, Lawrence Flynn, Christopher Gilbert, Ian Goodhead, Mareike Janiak, Laurent Marivaux, Michael Matschiner, Michèle Morgan, Christian Roos, and Erik Seiffert. Mareike Janiak helped write the code that generated Figures 1 and 2. Funding for this paper was provided by NERC Standard Grant “Rise of the continent of the monkeys” (NE/T000341/1).

