## Supplementary Online Material for "Total evidence tip-dating phylogeny of platyrrhine primates and 27 well-justified fossil calibrations for primate divergences"

**SOM Table S1**

Age priors of fossil taxa used in total evidence tip-dating analyses presented in this study. All extant taxa were given a fixed age prior of 0 Ma. As recommended by Püschel et al., (2020), age priors for fossil taxa were based on the total age range of the specimen(s) used for scoring the morphological matrix of Kay (2015; see also Kay et al., 2019), which we used as the source of our morphological data (see main text). As a result, the age ranges for a few taxa are broader than indicated in the "Fossil Calibrations" section of our main text, which only gives the age range of the oldest known specimen(s) for a particular taxon.

| Fossil taxon | Age prior (uniform, unless otherwise noted) | Justification |
|---|---|---|
| *Paralouatta* | 0 - 1.29 Ma | Described as late Quaternary based on associated fauna (Rivero and Arredondo, 1991). We use the second half of the Quaternary (Cohen et al., 2013) for this age prior. |
| *Xenothrix* | 0.001443 - 0.001511 Ma | Radiometric date reported by Cooke et al. (2017). |

| | | |
|---|---|---|
| *Antillothrix* | 0.004066 - 0.004458 Ma | Radiometric date reported by Rímoli (1977). |
| *Acrecebus* | 5.28 - 9.0 Ma | Huayquerian SALMA (Prevosti and Forasiepi, 2018, Table 1.1). See main text for more information on the age justification of this taxon. |
| *Proteropithecia* | 10.4 - 19.76 Ma | Known specimens of this taxon are derived from within and above the Pilcaniyeu Ignimbritic Member in the Collón Curá Formation (Kay et al., 1998a). The minimum bound for this age prior is based on a 10.6 +/- 0.2 Ma U-Pb date from the overlying Caleufu Formation (López et al., 2019); the maximum bound is based on a 19.04 +/- 0.72 Ma date from 40Ar/39Ar biotite dating of the basal ignimbrite in the Collón Curá Formation (Nivière et al., 2019). |
| *Neosaimiri* | 12.272 - 13.183 Ma | The type specimen of this taxon is from the Monkey Beds at La Venta, which correspond to the normal interval of Chron C5AA (Flynn et al., 1996; Kay and Madden, 1997). This interval spans from 13.183 to 13.032 Ma (Gradstein et al., 2012). *Neosaimiri* specimens described by Rosenberger et al. (1991) and Takai (1994), which were also used by Kay et |

| | | |
|---|---|---|
| | | al. (2019) for scoring of this taxon, are from the Masato site in the "lowest part of the Tatacoa Red Member”. According to Villarroael A. (1996), the Tatacoa Red Member is equivalent to the El Cardon Red Beds. Magnetostratigraphic information provided by Montes et al. (2021) indicates that the El Cardon Red Beds are younger than 12.829 Ma but older than 12.272 Ma. Therefore, the composite age range for this taxon is 12.272-13.183 Ma; this differs from the age used for the calibration of crown Cebidae (see main text) which uses the age estimate of the earliest occurrence of *Neosaimiri* (in the Monkey Beds) only. |
| *Nuciruptor* | 12.272-12.829 Ma | Age estimate of the El Cardon Red Beds at La Venta (see “*Neosaimiri*” above). |
| *“Aotus” dindensis* | 13.032 - 13.183 Ma | Age estimate of Monkey Beds at La Venta (see “*Neosaimiri*” above). |
| *Cebupithecia* | 13.032 - 13.183 Ma | Age estimate of Monkey Beds at La Venta (see “*Neosaimiri*” above). |
| *Mohanamico* | 13.032 - 13.183 Ma | Age estimate of Monkey Beds at La Venta (see “*Neosaimiri*” above). |

| | | |
|---|---|---|
| *Stirtonia* spp. | 13.032-13.608 Ma | The older species used by Kay (2015) for scoring purposes, namely *S. victoriae*, is from Duke Locality 28 at La Venta, within the Cerro Gordo Beds of the La Victoria Formation, which lies within Chron C5ABn (Montes et al., 2021); this spans from 13.608 to 13.363 Ma (Gradstein et al., 2012), with the latter date providing our minimum bound. The younger species used by Kay (2015) for scoring purposes, namely *S. tatacoensis*, is from the Monkey Beds and Fish Beds, which are older than 13.032 Ma (Montes et al., 2021). This age prior differs from the minimum age of the fossil calibrations of crown Platyrrhini and crown Atelidae above, as both *S. victoriae* and *S. tatacoensis* were used for scoring this genus in the morphological matrix used in our total evidence tip-dating analyses (Kay, 2015; Kay et al., 2019), whereas only *S. victoriae* (the older of the two species) was used for calibrating purposes. |
| *Lagonimico* | 13.183 - 13.608 Ma | Date for Duke/INGEOMINAS locality 90 (see age justifications of the Callitrichidae-Cebidae split in the main text for more information on this age estimate; Montes et al., 2021). |

| | | |
|---|---|---|
| *Homunculus* spp. | 16 - 18 Ma | All of the *Homunculus* specimens appear to be from Santacrucian sites on the Atlantic coastal plain (see Appendix 16.1 Kay et al., 2012), which are 18-16 Ma (Perkins et al., 2012). |
| *Parvimico* | 16.4 - 19.6 Ma | Radiometric date reported by Kay et al. (2019). |
| *Carlocebus* spp. | 16.84 - 18.01 Ma | *Carlocebus* spp. are from the Pinturas Formation. According to Perkins et al. (2012), the overlying Toba Blanca tuff is 16.89 +/- 0.05 Ma, giving a minimum age of 16.84 Ma, and a date from near the base of the formation at the Estancia El Carmen is 17.99 +/- 0.02 Ma, which we use as a maximum of 18.01 Ma. |
| *Soriacebus* spp. | 16.84 - 18.01 Ma | *Soriacebus* is from the Pinturas Formation (see "*Carlocebus* spp." above) |
| *Panamacebus* | 18.748 - 21.1 Ma | The overlying Centenario Fauna is within C5Er (MacFadden et al., 2014), which is 18.748-18.524 Ma according to Gradstein et al. (2012); 18.748 Ma is therefore the minimum bound for this age prior. The maximum bound is the maximum age of the radiometric date underlying the fossil deposit, as |

| | | |
|---|---|---|
| | | presented by Bloch et al. (2016). |
| *Chilecebus* | 19.82 - 20.36 Ma | Radiometric date reported by Flynn et al. (1995). |
| *Mazzonicebus* | 20.040 - 21.083 Ma | Material came from the Colhue-Huapi West locality, the Lower Fossil Zone (LFZ) of the Colhue-Huapi Member, Sarmiento Formation at Gran Barranca (Kay, 2010). The LFZ corresponds to Chron C6An.1n (Ré et al., 2010), which, following Gradstein et al. (2012) is 20.040-20.213 Ma. However, according to Dunn et al. (2013), it spans C6An.2r to C6An.2n, which is 21.083 to 20.439 Ma following Gradstein et al. (2012). We use the maximum possible age range for this age estimates here. |
| *Dolichocebus* | 20.1 - 21.0 Ma | Material comes from the Trelew Member of the Sarmiento Formation which belongs to the Colhuehuapian SALMA (Kay et al., 2008). We follow the age estimate for the Colhuehuapian SALMA of Dunn et al. (2013). |
| *Tremacebus* | 20.1 - 21.0 Ma | Materials come from Colhuehuapian aged deposits at approximately 12 km southwest of Cerro Sacanana, in north central Chubut (Hershkovitz, 1974). |

| | | |
|---|---|---|
| | | We follow the age estimate for the Colhuehuapian SALMA of Dunn et al. (2013). |
| *Canaanimico* | 23 - 26.63 Ma | The minimum bound is provided by the minimum age of the Deseadan SALMA (Dunn et al., 2013). The maximum bound is based on the underlying, dated tuff that is ~5m below the fossils (Marivaux et al., 2016). |
| *Branisella* | 25.264 - 25.304 Ma | Kay et al. (1998b) correlated the "Branisella Zone/Level" (= Unit 5) at Salla to C8n.lr; according to Gradstein e al. (2012), this is 25.304-25.264 Ma. |
| *Perupithecus* | 29.52 - 29.68 Ma | Radiometric date reported by Campbell et al. (2021). |
| *Aegyptopithecus* | 28.2 - 33.4 Ma | Most specimens of *Aegyptopithecus* appear to be from "Quarry M", which is one of the youngest sites in the Gebel Qatrani Formation; however, the age of the Gebel Qatrani Formation is somewhat controversial (see Van Couvering and Delson, 2020), and so we assign an age representing the entire of the "Qatranian" (sensu Van Couvering and Delson, 2020), which is 33.4-28.2 Ma. |

| | | |
|---|---|---|
| *Apidium* | 28.2 - 33.4 Ma | See "*Aegyptopithecus*" above. |
| *Simonsius* | 28.2 - 33.4 Ma | See "*Aegyptopithecus*" above. |
| *Catopithecus* | 33.4 Ma (fixed) | This taxon is from Quarry L-41 in the Fayum Depression, Egypt; the fixed age estimate for this site follows Van Couvering and Delson (2020). |
| *Proteopithecus* | 33.4 Ma (fixed) | See "*Catopithecus*" above. |
